# A scalable human neuromuscular organoid platform enables lineage-specific analysis of drug responses in spinal muscular atrophy

**DOI:** 10.64898/2026.08.23.745904

**Authors:** Ines Lahmann, Angélica García-Pérez, Ismail Amr El-Shimy, Inês Afonso Martins, Lan Vi Ngoc Nguyen, Chrysanthi-Maria Moysidou, Noelle Findeisen, Ina-Maria Rudolph, Christina Bukas, Donatella Cea, Gary J. Bassell, Wilfried Rossoll, Marie Piraud, Sebastian Diecke, Mina Gouti

## Abstract

Scalable human models that capture interactions between distinct tissues remain limited, constraining mechanistic insight and therapeutic prediction. Here, we established a scalable, automation-compatible human neuromuscular organoid (NMO) platform that enables integrated analysis of neuronal and muscle lineages in spinal muscular atrophy (SMA). Patient-derived NMOs reproducibly self-organise into spinal cord and skeletal muscle compartments and form functional neuromuscular circuits. SMA NMOs recapitulate early disease features, including reduced survival motor neuron (SMN) protein levels and impaired neuromuscular junction (NMJ) maturation. Single-nucleus RNA sequencing identifies lineage-specific transcriptional changes across neuronal and muscle compartments preceding functional deficits. Using this platform, we compared two clinically relevant SMN2 splicing modulators and observed distinct, cell-type-dependent responses. While both compounds increased SMN levels and NMJ number, only one enhanced myofiber growth and improved contractile function. These findings highlight muscle maturation, rather than NMJ number alone, as a key determinant of functional recovery and establish NMOs as a scalable system for studying cell-type-specific therapeutic responses.

## Introduction

Many human diseases arise from failures of interactions between distinct but interdependent tissues. Yet, replicating these multicellular dynamics *in vitro* remains a fundamental challenge. Conventional models typically focus on isolated cell types or single tissue systems, overlooking emergent mechanisms that depend on reciprocal signalling and co-maturation across lineages. This limitation has hindered efforts to study the integrated pathophysiology of complex disorders, such as neuromuscular diseases^1,2^, and reflects a broader challenge of connecting cellular-scale molecular data to emergent tissue-level function^3,4^, thereby constraining mechanistic understanding and therapeutic development. Human neuromuscular organoids (NMOs) address this challenge by self-organising from human pluripotent stem cell (hPSC)-derived neuromesodermal progenitors (NMPs)^5,6^ into coordinated neural and mesodermal compartments that give rise to functional neuromuscular junctions (NMJs)^7–9^. By recapitulating *in vivo* developmental processes^10^, NMOs establish bidirectional neural-muscle connectivity and provide a quantifiable readout of human circuit activity via spontaneous muscle contraction^7^.

We applied this approach to spinal muscular atrophy (SMA), one of the most common genetic causes of infant mortality^11^. SMA is primarily caused by homozygous deletions or mutations in the Survival of Motor Neuron 1 (*SMN1*) gene, resulting in reduced levels of the ubiquitously expressed SMN protein^11–13^. The establishment of the first SMA patient-derived induced pluripotent stem cell (iPSC) line provided a human platform for modelling motor neuron pathology in vitro^14^, and laid the foundation for studies extending beyond the neural lineage^8,15^. Disease severity inversely correlates with the number of *SMN2* copies, a nearly identical paralog of *SMN1* that predominantly produces a truncated, non-functional SMN protein due to exon 7 skipping during pre-mRNA splicing^16,17^. Although SMA is considered a monogenic disease, it exhibits substantial clinical heterogeneity even among patients with identical *SMN1* and *SMN2* copy numbers, suggesting that additional modifiers influence disease progression^18,19^. Early studies demonstrated that SMA patient myoblasts exhibit fusion defects and impaired acetylcholine receptor clustering^20^, and accumulating evidence indicates that skeletal muscle contributes independently to SMA pathogenesis^15^. Together, these observations support a model in which SMA arises from impaired neuromuscular system integration rather than strictly autonomous neuron dysfunction. SMN is known to play central roles in RNA processing and ribonucleoprotein assembly, and its deficiency has been linked to widespread defects in RNA metabolism and cellular homeostasis^21,22^, while additional studies have highlighted mitochondrial dysfunction as a key contributor to motor neuron vulnerability in SMA^23^.

Over the past decade, therapeutic strategies targeting SMN restoration have transformed SMA treatment. These include antisense oligonucleotides and gene-replacement approaches, as well as small molecules that modulate SMN2 splicing^18,24,25^. Among these, orally available splicing modifiers such as Risdiplam have shown substantial clinical benefit by promoting exon 7 inclusion and increasing full-length SMN protein levels^25,26^. Branaplam, another SMN2 splicing modulator^27^, also demonstrated therapeutic potential, but was discontinued due to safety concerns^28^. However, variability in therapeutic response remains a key challenge, and emerging evidence suggests that different splicing modulators may elicit distinct downstream transcriptional effects, raising the possibility of lineage-specific mechanisms of action^29,30^ as well as patient-specific responses^25^. Recent spinal cord organoid models have revealed early neurodevelopmental defects and neuromesodermal lineage imbalances in SMA, suggesting that disease pathogenesis may originate during development^31^; however, the absence of functional neuromuscular integration in these systems limits their ability to link early defects to circuit-level dysfunction. These findings highlight the urgent need for physiologically relevant, multi-lineage, patient-specific human models capable of resolving cell-type-specific drug mechanisms and linking them to functional outcomes^1,32^. Here, we focus on small-molecule SMN2 splicing modulators^26,27^, which provide a tractable system to dissect drug-specific and cell-type-specific mechanisms in human neuromuscular tissues.

To enable systematic and scalable interrogation of SMA pathology, we developed a robot-assisted platform for the reproducible generation of NMOs from healthy and patient-derived iPSCs. This automated workflow reduced operator-dependent variability while preserving disease-relevant phenotypes, allowing the parallel production of large numbers of NMOs suitable for downstream functional and molecular analyses. We generated NMOs from three SMA type 1 patients, which recapitulate coordinated spinal cord and skeletal muscle development and enable the study of neuromuscular circuit formation in a human context. SMA NMOs exhibited early, multi-lineage pathology involving neurons, muscle fibers and NMJs, consistent with impaired neuromuscular circuit integrity.

To dissect the molecular basis of these phenotypes, we performed single-nucleus RNA sequencing (snRNA-seq)^33,34^, which revealed distinct, lineage-specific transcriptional alterations, including synaptic and metabolic dysfunction in neurons and stress-associated maturation and apoptosis signatures in muscle. These changes converged on a shared phenotype of impaired NMJ stability. Within this framework, we investigated the effects of clinically relevant SMN2 splicing modulators and observed that, although SMN restoration improved NMJ number, downstream transcriptional and functional responses differed across cell types. Notably, enhancement of muscle-intrinsic programs was associated with improved neuromuscular function beyond SMN restoration alone. Together, these findings support a model in which SMA involves impaired stabilization of a multi-lineage neuromuscular circuit and establish NMOs as a scalable human platform for resolving lineage-specific disease mechanisms and therapeutic responses.

## Results

### A scalable and automated human neuromuscular organoid platform

To enable reproducible, scalable, and systematic interrogation of patient-specific neuromuscular phenotypes, we developed a robot-assisted differentiation workflow based on a Biomek i7 platform and combined it with the large-scale culture and cryopreservation of neuromesodermal progenitors (NMPs). This modular system allows production of NMOs from multiple pluripotent stem cell (PSC) lines while reducing operator-dependent variability (Fig.1a, Extended Data Fig.1a-d).

For the automated generation of NMOs, iPSCs were maintained on large-scale cell culture dishes (OneWell) and the automated system was used to dissociate, count, and seed them to generate NMPs. Three days after plating, these cultures showed typical NMP colony formation and the expression of the NMP markers (*TBXT*, *SOX2*) and *TBX6* (Fig. 1b). NMPs were dissociated using the automated system and cryopreserved or seeded in 96-plates for downstream generation of NMOs. In a single automated run, three OneWell plates can be processed in parallel, yielding approximately 60 million NMPs and enabling large-scale production of NMOs. Under standard conditions, approximately 900,000 NMPs are seeded per 96-well plate, corresponding to 9,000 cells per organoid. In routine experiments, we generated 1-6 96-well plates per run (96-576 NMOs), with the capacity to scale to several thousand organoids if required.

**Figure 1.**
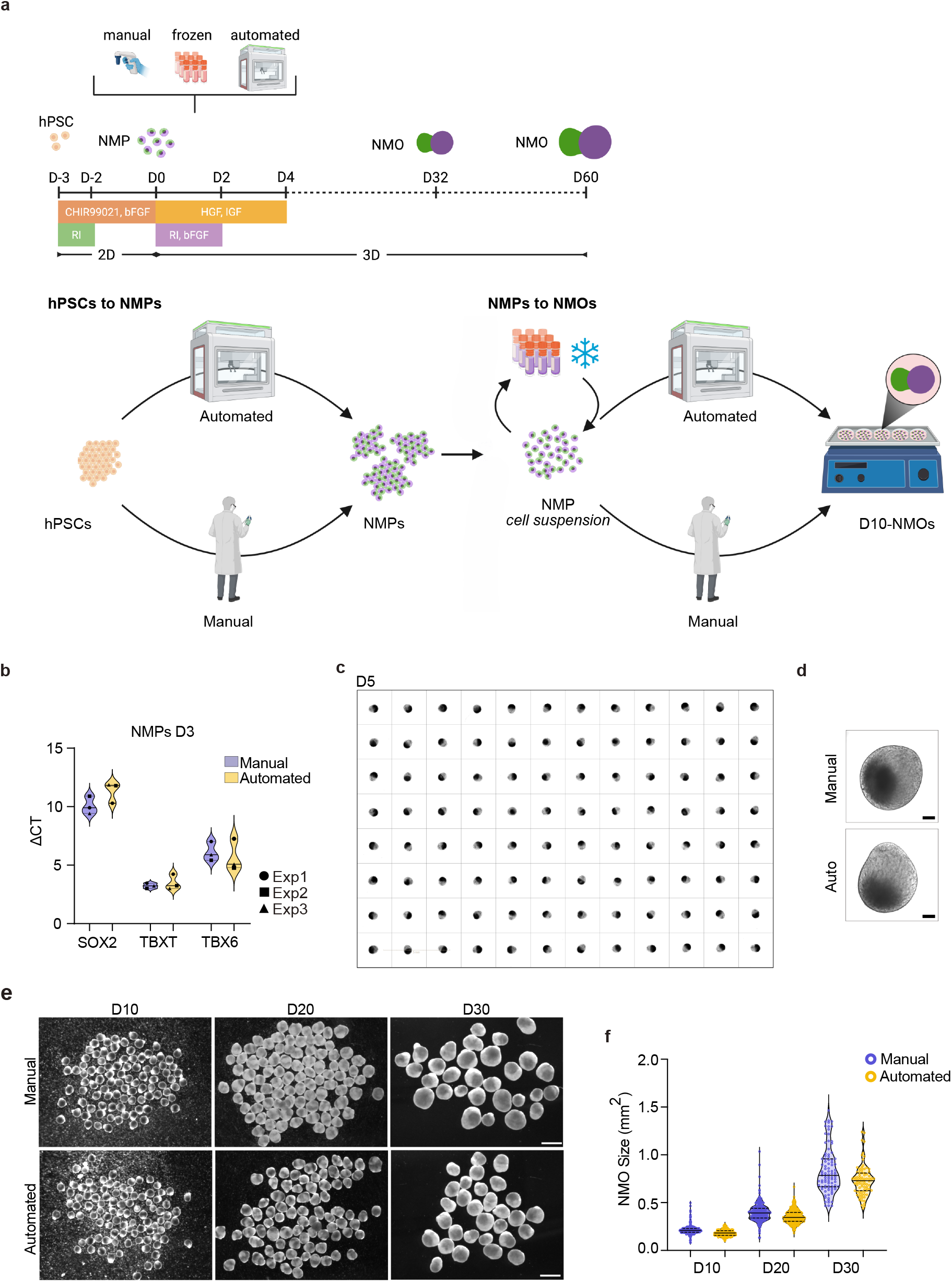
Automated workflows support scalable generation of NMOs. (a) Schematic outline of manual and automated workflows from human iPSCs to NMPs, and from NMPs to day 10 NMOs, including an optional cryopreservation step.| (b) RT-qPCR analysis of NMP markers (*SOX2*, *TBXT*, *TBX6*) in control NMPs derived from manual and automated protocols. (c) Brightfield images of day 5 control NMOs generated from frozen NMPs in a 96-well format. (d) Representative brightfield images of day 5 control NMOs derived from manual and automated workflows. (e) Representative brightfield images of day 10, 20 and 30 control NMOs generated from manual, frozen, or automated workflows. (f) Organoid size over time (days 10, 20, and 30) in control NMOs derived from manual and automated protocols. (g) Representative brightfield images of day 60 control NMOs generated from manual or automated workflows. (h) Representative immunofluorescence images of day 60 NMOs stained for neurons (TUBB3, yellow), muscle (MYH1/2, magenta), and acetylcholine receptor clusters (αBTX, cyan). Arrowheads indicate αBTX clusters. (i) Quantification of NMJ number (αBTX⁺ clusters) per field of view (FOV) in control NMOs generated using manual and automated workflows. Scale bars: (c) 5 mm, (d) 500 µm, (e, g) 1 mm, (h) 250 µm and 25 µm. Data in b, f and i are presented as violin plots showing the median, quartiles and individual data points. Each dot represents one NMO in b and f. Different symbols denote independent experiments. For b and i, N = 3 independent experiments and n = 3 per condition. For f, N = 1 and n = 96 per condition and time point. In i, each dot represents one field of view per NMO; different symbols indicate independent NMOs.

To assess robustness across workflows, we compared NMOs generated manually, from cryopreserved NMPs, and via automated differentiation (Fig.1a). Organoid formation was first assessed by high-content imaging at day 1 and day 5. All conditions produced elongated NMOs across wells at day 5 (Fig.1c and Extended Data Fig.1a,c). Morphological analysis showed comparable size and elongation across conditions, consistent with proper early patterning (Fig.1d and Extended Data Fig.1e,f). Notably, automated NMOs displayed reduced variability in size at both day 1 and day 5 compared to manual workflows performed by three independent operators (User 1-3), each generating three 96-well plates in parallel (Extended Data Fig. 1a-f), indicating improved size reproducibility. Growth dynamics between day 10 and day 30 were similar across workflows (Fig.1e,f), indicating that automation preserves the kinetics of organoid formation. By day 60, both manual and automated workflows yielded well-compartmentalised NMOs with spatially segregated neural and muscle domains that organise into NMJ structures consistent with the expected architecture of neuromuscular organoids^7^ (Fig.1g,h and Extended Data Fig.1g,h). Together, these results demonstrate that automated differentiation and NMP banking enable the reproducible and scalable generation of self-organising NMOs with preserved neuromuscular architecture and NMJ formation, providing a foundation for downstream disease modelling and therapeutic testing.

### Patient-derived NMOs reveal early multi-lineage SMA pathology

We next investigated whether patient-derived NMOs generated under standard culture conditions capture disease-relevant phenotypes across interacting neuromuscular lineages in SMA. To establish a human model of SMA, we generated neuromuscular organoids (NMOs)^7^ from three SMA type 1 patient-derived iPSC lines with varying disease severity (Pt1-Pt3; Fig.2a, and Extended Data Fig.2a). Given the complexity of the SMN locus, we included independent healthy control pluripotent stem cell (PSC) lines, derived from both embryonic and induced sources, which showed consistent phenotypes across lines, as previously reported for NMOs^7^ (Extended Data Fig.2a).

All three SMA lines reproducibly formed polarised NMOs with distinct neural and mesodermal compartments, comparable to controls (Fig.2a,b and Extended Data Fig.2b). At an early developmental stage (day 20), organoids exhibited the expected segregation of neural and mesodermal compartments, along with reduced SMN protein levels in SMA lines (Extended Data Fig.3a, b). By day 50, SMA-NMOs displayed overall morphology and tissue organisation similar to controls, including well-defined compartmentalisation and broadly preserved tissue proportions (Extended Data Fig.3c-e).

To assess neuromuscular development, we quantified postsynaptic acetylcholine receptor (AChR) clusters using α-bungarotoxin (αBTX) staining at day 30, corresponding to early NMJ formation and at day 60, when NMJs undergo maturation and support spontaneous motor neuron-driven contractions^7^. AChR clusters were normalised to myofiber number and are hereafter referred to as neuromuscular junctions (NMJs), representing postsynaptic specialisations associated with neuromuscular contacts. Across three independent differentiations of the three SMA patient lines, SMA NMOs exhibited a robust and significant reduction in NMJ number at day 60 compared to controls (Fig. 2c), consistent with impaired NMJ formation^35^. This phenotype was preserved across NMOs generated manually, from cryopreserved NMPs, versus the automated workflow (Extended Data Fig.4e).

**Figure 2.**
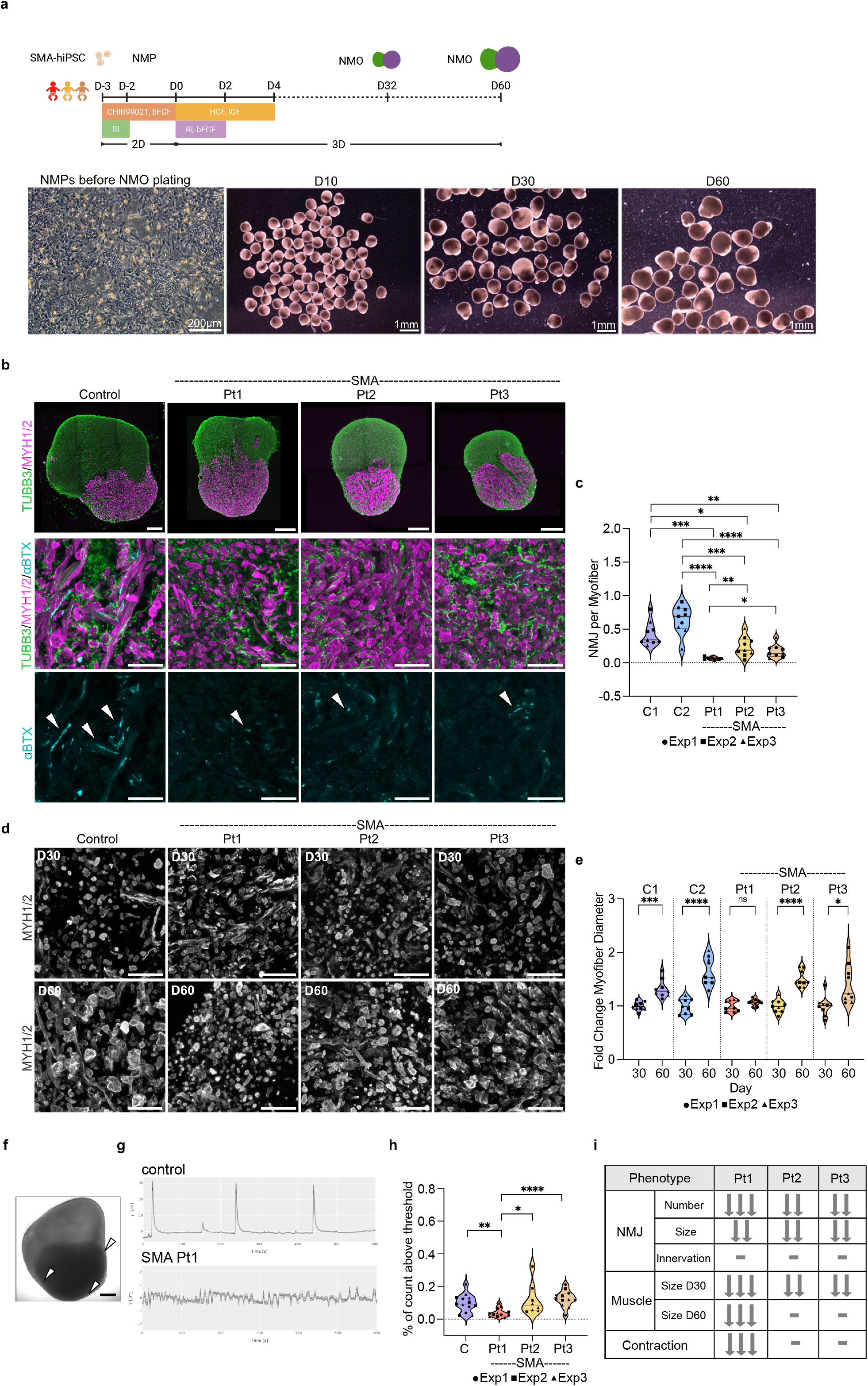
Efficient generation of iPSC-derived SMA NMOs reveals NMJ deficits and impaired myofiber maturation. (a) Schematic illustration of the differentiation protocol for generating NMOs. Human SMA iPSCs were directed toward NMPs and subsequently aggregated to form 3D NMOs. Representative brightfield images show day 3 NMPs and SMA NMO Pt1 organoid morphology at days 10, 30, and 60. (b) Representative immunofluorescence images of day 60 NMOs control and SMA NMOs from three patient lines (Pt1–Pt3), stained for neurons (TUBB3, green), muscle (MYH1/2, magenta), and acetylcholine receptor clusters (α-bungarotoxin, αBTX; cyan). Arrowheads indicate αBTX clusters. (c) Quantification of NMJs as αBTX⁺ clusters per myofiber at day 60 NMOs. (d) Immunofluorescence images of muscle regions of NMOs at day 30 and day 60 from control and SMA patient lines stained for MYH1/2. (e) Fold change in myofiber diameter between day 30 and day 60. (f) Representative brightfield image of an NMO before contraction recording. Arrowheads mark the three regions selected for data acquisition. (g) Representative Time Series plots of spontaneous contractions during time-lapse recordings from control and SMA Pt1 NMOs. (h) Quantification of NMO contractile activity, expressed as the percentage of active points above threshold. (i) Qualitative summary of the NMJ, skeletal muscle and functional phenotype in SMA patient-derived NMOs compared to control. Scale bars: (a) 200 µm and 1 mm, (c) 250 µm and 50 µm, (d) 50 µm. Data are presented as violin plots displaying the median, quartiles, and individual data points. Each dot represents one NMO (c, e, h). Different symbols indicate independent experiments, with N = 3 and n = 3 for all conditions in panels c, e, and h. Statistical significance was assessed using a two-sided Welch’s t-test to account for unequal variances between groups (*P < 0.05; **P < 0.01; ***P < 0.001; ****P < 0.0001).

Morphometric analysis showed that NMJ size was comparable at day 30, but failed to increase in SMA NMOs by day 60, resulting in significantly smaller NMJs (Extended Data Fig.4a-c), consistent with impaired synaptic stabilisation rather than defective initial formation^35^. Despite their reduced size, residual AChR clusters in SMA organoids remained innervated, as indicated by apposition to neuronal projections (Extended Data Fig.4d), suggesting that neuromuscular contacts are formed but do not fully mature. These phenotypes were consistently observed across all three SMA lines, supporting a robust and disease-relevant neuromuscular phenotype.

We next examined whether NMJ defects were accompanied by alterations in the muscle compartment. While overall tissue organisation was preserved, SMA NMOs displayed an increased number of cleaved Caspase-3 (cCasp3) positive myofibers, indicating increased vulnerability of the muscle compartment, whereas motor neuron apoptosis (ChAT^+^/cCasp3^+^ cells) remained limited at this stage (Extended Data Fig.5a-d). Consistent with this, SMA NMOs exhibited an apparent increase in myofiber number at day 60 compared to controls, accompanied by a reduction in myofiber diameter across all three patient lines (Extended Data Fig.4f-h). This reduction was already apparent at day 30 and became more pronounced by day 60, particularly in the Pt1 line.

Analysis of myofiber growth between day 30 and day 60 revealed both shared and patient-specific defects. Although myofiber size increased over time in control NMOs as well as Pt2 and Pt3 lines, these fibres remained smaller and more numerous than in control. In contrast, Pt1 myofibers failed to increase in size (Fig.2d,e and Extended Data Fig. 4f-i), indicating a more severe impairment in muscle maturation in the most affected line^20^.

To assess circuit-level function, we quantified spontaneous contraction in NMOs from all patient lines. SMA Pt1 NMOs exhibited a profound reduction in Contraction power (μm^2^/s), reflecting the magnitude of tissue displacement over time and contraction frequency (% of count above threshold), whereas Pt2 and Pt3 NMOs retained measurable contractile activity (Fig.2f,g,h; Extended Data Fig 4j) (Supplementary Movies 1-4). Pharmacological blockade with curare abolished contractions across all conditions (Supplementary Movies 5-8), confirming that contractile activity depends on motor neuron-driven neuromuscular transmission.

Together, these findings demonstrate that SMA NMOs recapitulate key features of early multi-lineage pathology, characterised by preserved tissue architecture but impaired NMJ maturation and stabilisation. These defects are accompanied by muscle-specific vulnerability and patient-dependent functional impairment, supporting a model in which neuromuscular dysfunction arises from combined deficits in synaptic maturation and muscle growth.

### Single-nucleus RNA-seq reveals lineage-specific transcriptional vulnerabilities in SMA NMOs

To uncover the molecular basis of the multi-lineage phenotypes observed in SMA NMOs, we performed single-nucleus RNA sequencing (snRNA-seq) at day 30, a stage preceding the defects in NMJ maturation observed at day 60 (Fig.3 and Extended Data Fig.6a-c). We profiled an average of ∼3,800 nuclei isolated from three independent NMOs per line, capturing both control and SMA samples. Unsupervised clustering resolved 18 transcriptionally distinct cell clusters at day 30 (Fig.3a), with clear segregation of neural and mesodermal lineages. These populations were consistently detected across control and SMA NMOs, with largely preserved relative abundances (Extended Data Fig.6a,b), indicating that SMN deficiency does not substantially alter cell-type composition at this stage. The individual clusters expressed gene signatures associated with specific cellular identities, including neural progenitors, spinal neurons, glia, Schwann cells, muscle progenitors, skeletal muscle cells and fibroblasts (Extended Data Fig. 6b,c).

**Figure 3:**
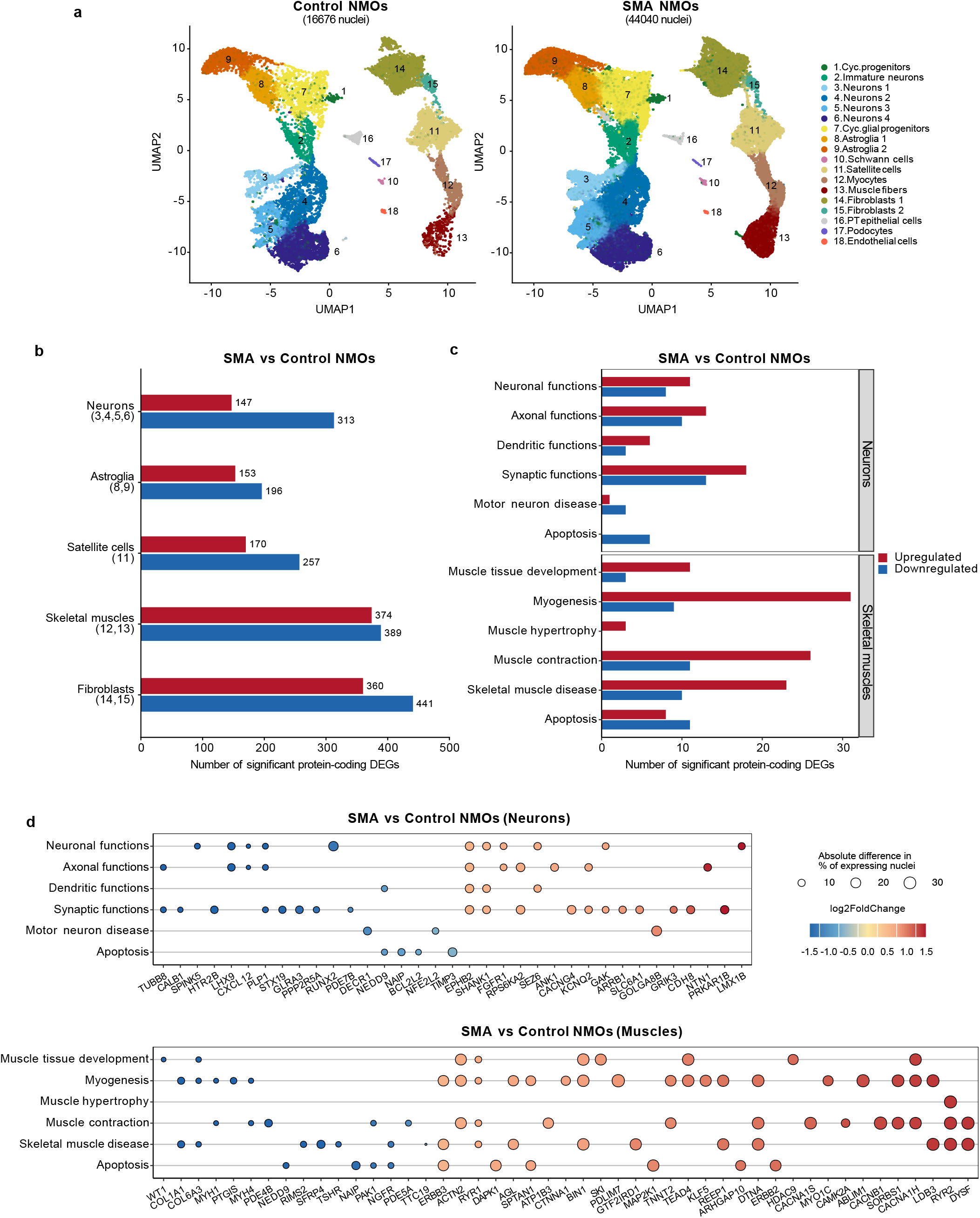
Single-nucleus transcriptomic analysis reveals early lineage-specific SMA-associated transcriptional changes in neurons and skeletal muscle at day 30 NMOs. (a) UMAP representations of SMA (n = 44,040) and control (n = 16,676) NMO samples showing 18 transcriptionally distinct nuclear clusters of both neuronal (10 clusters) and mesodermal (8 clusters) lineages. (b) Differential gene expression analysis (DESeq2) comparing SMA and control NMOs across major cell types, including merged neuronal clusters (3-6), astroglia clusters (8-9), satellite cells (11), skeletal muscle clusters (12-13) and fibroblasts (14-15). Bars indicate the number of significantly up and downregulated protein-coding differentially expressed genes (DEGs) per cell type. Gene significance cutoffs are set to an absolute fold change > 1.5 and an adjusted P value < 0.05. (c) Functional categorisation of DEGs across neuronal and muscle populations, highlighting enrichment of biological processes related to synaptic signalling, neuronal connectivity, and cytoskeletal organisation in neurons and muscle contraction, myogenesis and structural remodelling pathways in skeletal muscle. (d) Representative DEGs illustrating lineage-specific transcriptional programs in neurons and skeletal muscle, grouped by cellular functional categories. Dot color indicates log_2_ fold change (SMA vs control) and dot size represents the absolute difference in the percentage of gene-expressing nuclei between conditions.

We next performed differential gene expression analysis to identify lineage-specific transcriptional changes associated with SMN deficiency (Fig.3b,c and Supplementary Table 1). SMA NMOs exhibited distinct transcriptional changes across the neural and mesodermal cell types, which were consistently detected across the three patient lines, indicating a reproducible response to SMN deficiency (Fig.3b,c). In neuronal populations, SMA NMOs showed transcriptional changes in gene programs associated with synaptic function, neuronal excitability and connectivity. Upregulated genes included neurotransmitter receptors and synaptic regulators (GRIK3, SLC6A1), ion channels (KCNQ2, CACNG4) and adhesion molecules (CDH8), indicating coordinated alterations in synaptic and excitability-related pathways^36^. In parallel, genes associated with axon guidance and neuronal connectivity, including *NTN1* and *EPHB2*, were upregulated, consistent with a shift in connectivity-related transcriptional programs^35,37,38^. These changes are consistent with a transcriptional state characterised by altered synaptic signalling and connectivity-related gene expression programs (Fig.3c,d).

In contrast, skeletal muscle populations exhibited a distinct transcriptional response involving genes associated with myogenesis, myofibril assembly, and muscle contraction (Fig.3c,d). Multiple genes involved in calcium handling, structural integrity and membrane repair, including *RYR1, CACNB1* and *DYSF*, were upregulated^39^. Additional changes were observed in extracellular matrix components (e.g., *COL1A1*, *COL6A3*) and cytoskeletal regulators (Fig.3d), indicating alterations in structural and remodelling-related pathways. Consistent with these gene-level changes, gene set enrichment analysis showed enrichment of pathways related to myogenesis, myofibril assembly, actomyosin organisation, and muscle contraction, supporting broad alterations in muscle differentiation and functional programs^40,41^ (Fig.3c,d and Supplementary Table 2). Together, these findings indicate an altered muscle transcriptional state in SMA NMOs, characterised by concurrent changes in muscle differentiation, structural organisation, and remodelling-associated pathways. Notably, these lineage-specific transcriptional changes occur in the absence of detectable differences in cell-type composition and precede the structural and functional NMJ defects observed at later stages, suggesting that early molecular perturbations may contribute to subsequent phenotypic outcomes (Fig.3a-d and Extended Data Fig.6a-c).

Together, these findings demonstrate that SMN deficiency induces early, lineage-specific transcriptional changes in both neuronal and muscle compartments. These distinct but concurrent alterations across neuromuscular lineages provide a molecular framework that may contribute to the subsequent emergence of neuromuscular dysfunction in SMA NMOs, linking early transcriptional perturbations to later structural and functional deficits.

### Lineage-specific effects of SMN2 splicing modulators in SMA NMOs

Given the severe phenotype observed in SMA Pt1 NMOs, including impaired neuromuscular function, we evaluated the effects of two clinically relevant SMN2 splicing modulators, Risdiplam and Branaplam (Fig.4a). Dose-response titrations identified concentrations compatible with organoid viability and structural integrity (250 nM Risdiplam and 40 nM Branaplam) (Fig.4a,b). At these concentrations, both compounds increased SMN protein levels across all three SMA patient lines (Fig.4b,c and Extended Data Fig.7a-c), and these conditions were used for all subsequent analyses.

**Figure 4.**
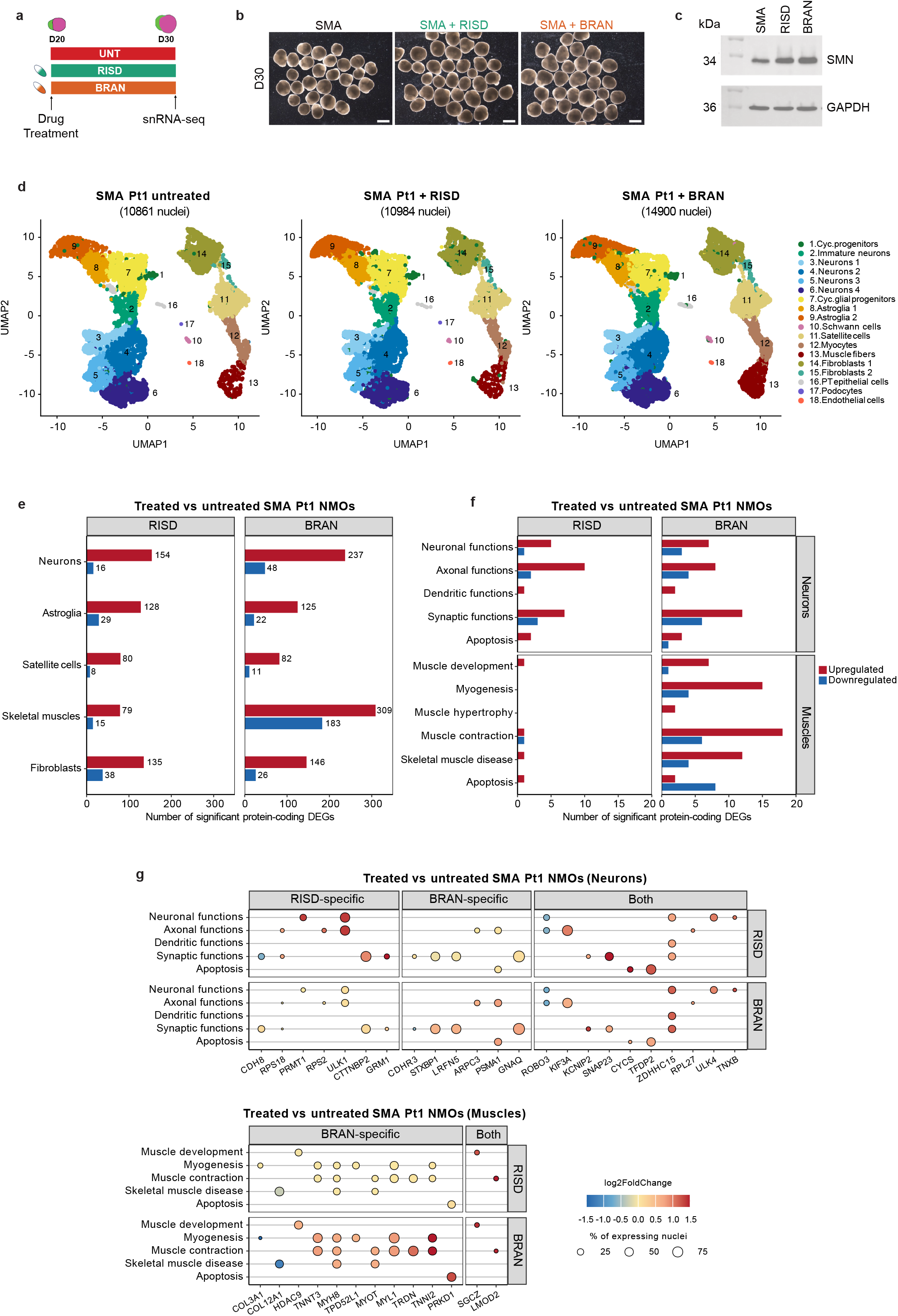
Risdiplam and Branaplam treatment induce distinct lineage-specific transcriptional responses in SMA NMOs. (a) Schematic of drug treatment and snRNA-seq timeline. SMA Pt1-derived NMOs were treated with Risdiplam (250nM) or Branaplam (40nM) from day 20 to day 30 and processed for snRNA-seq. (b) Representative brightfield images of day 30 NMOs under each condition (untreated, Risdiplam, Branaplam). (c) Western blot showing SMN protein (34kDa) levels in treated NMOs compared with untreated SMA Pt1 NMOs. GAPDH (36kDa) was used as a loading control. (d) UMAP representations of untreated (n = 10,861), Risdiplam-treated (n = 10984) and Branaplam-treated (n = 14,900) SMA Pt1 NMO samples showing comparable cell type composition across conditions. (e) Differential gene expression analysis using DESeq2 in treated vs untreated NMOs shows a more significant impact of Branaplam on NMO transcriptome across different cell types in terms of numbers of significant up/downregulated protein-coding DEGs. Gene significance cutoffs are set to an absolute fold change > 1.5 and a Benjamini-Hochberg-adjusted P value < 0.05. (f) Functional categorisation of DEGs associated with Risdiplam and Branaplam treatment, highlighting differences in affected biological processes across cell types including skeletal muscle. (g) Representative examples of interesting DEGs that are deregulated specifically by Risdiplam, Branaplam or both and the cellular functions they control are shown in neurons and muscles. Log_2_ fold change in gene expression in Risdiplam/Branaplam-treated vs untreated are represented as dot color and the percentage of gene-expressing nuclei in each treatment group as dot size. Scale bars: (b) 1 mm. RISD = Risdiplam, BRAN = Branaplam

To assess lineage-specific responses to SMN restoration, SMA NMOs were treated from day 20 to day 30 and subjected to single-nucleus RNA sequencing. Clustering analysis revealed comparable cellular composition across untreated and treated conditions across three biological replicates (Fig.4d), indicating that neither compound substantially alters lineage representation. Differential expression analysis demonstrated that both compounds induce transcriptional changes across multiple cell types; however, Branaplam elicited a broader and more pronounced response across both neuronal and muscle lineages (Fig.4e,f and Extended Data Fig.7d).

Comparison of drug-induced transcriptional programs revealed divergent lineage-specific effects despite a shared upstream mechanism of SMN restoration. In neuronal populations, both compounds modulated genes associated with synaptic signalling and neuronal function, with partially overlapping targets (Fig.4g). In contrast, skeletal muscle populations exhibited a distinct response. Branaplam selectively upregulated genes associated with myogenic differentiation and contractile maturation, including *TNNI2* and *MYL1*, which are linked to fast-twitch muscle fibre identity and contractile function (Fig.4g). Risdiplam did not induce activation of these myogenic programs, indicating that SMN restoration alone is insufficient to drive muscle maturation pathways under these conditions (Fig.4g).

To further dissect the nature of these lineage-specific responses, we classified treatment responsive genes as either reversed, reflecting restoration toward control expression levels, or potentiated, representing genes whose SMA associated dysregulation was further enhanced in the same direction upon treatment (Extended Data Fig.7e). Both compounds modulated disease-associated transcriptional signatures, with Branaplam exerting a broader and more pronounced effect (Extended Data Fig.7e,f). Reversed genes in neurons were enriched for synaptic and axonal functions, whereas in muscle cells, treatment-responsive genes included both reversed and potentiated genes associated with contractile and structural programs (Extended Data Fig.7g).

Skeletal muscle cells displayed a stronger transcriptional response to Branaplam than to Risdiplam, marked by potentiation of genes associated with muscle growth and contractile function, consistent with its broader transcriptional impact and activation of muscle intrinsic programs. A subset of these potentiated genes corresponded to pathways that were already upregulated in SMA NMOs, consistent with a potential compensatory response to muscle pathology, and were further enhanced upon Branaplam treatment (Extended Data Fig.7g). In parallel, comparison with previously reported off-targets of SMN2 splicing modulators^29^, defined as genes with altered splicing and expression upon treatment, revealed similar compound-specific transcriptional changes across multiple cell types, consistent with prior reports (Extended Data Fig. 7h).

Given these distinct transcriptional and mechanistic profiles, we next asked whether they translate into differential functional recovery at later stages of organoid maturation (day 60) (Fig.5a). Both compounds increased NMJ number relative to untreated SMA NMOs, with Risdiplam inducing an approximately two-fold increase and Branaplam a stronger (∼4-fold) increase (Fig.5b, c and Extended Data Fig.8a, b). However, these structural improvements did not translate equally into functional recovery. Risdiplam-treated NMOs showed minimal improvement in contractile activity, whereas Branaplam treatment partially restored contraction signal energy (Fig.5d and Extended Data Fig.8c,d; Supplementary movies S9-S11). Consistent with the lineage-specific transcriptional responses, only Branaplam significantly increased myofiber diameter (Fig.5e-g), indicating enhanced muscle maturation. These findings demonstrate that restoration of NMJ number alone may not be sufficient to recover neuromuscular function and instead identify muscle maturation as a key determinant of functional recovery.

**Figure 5.**
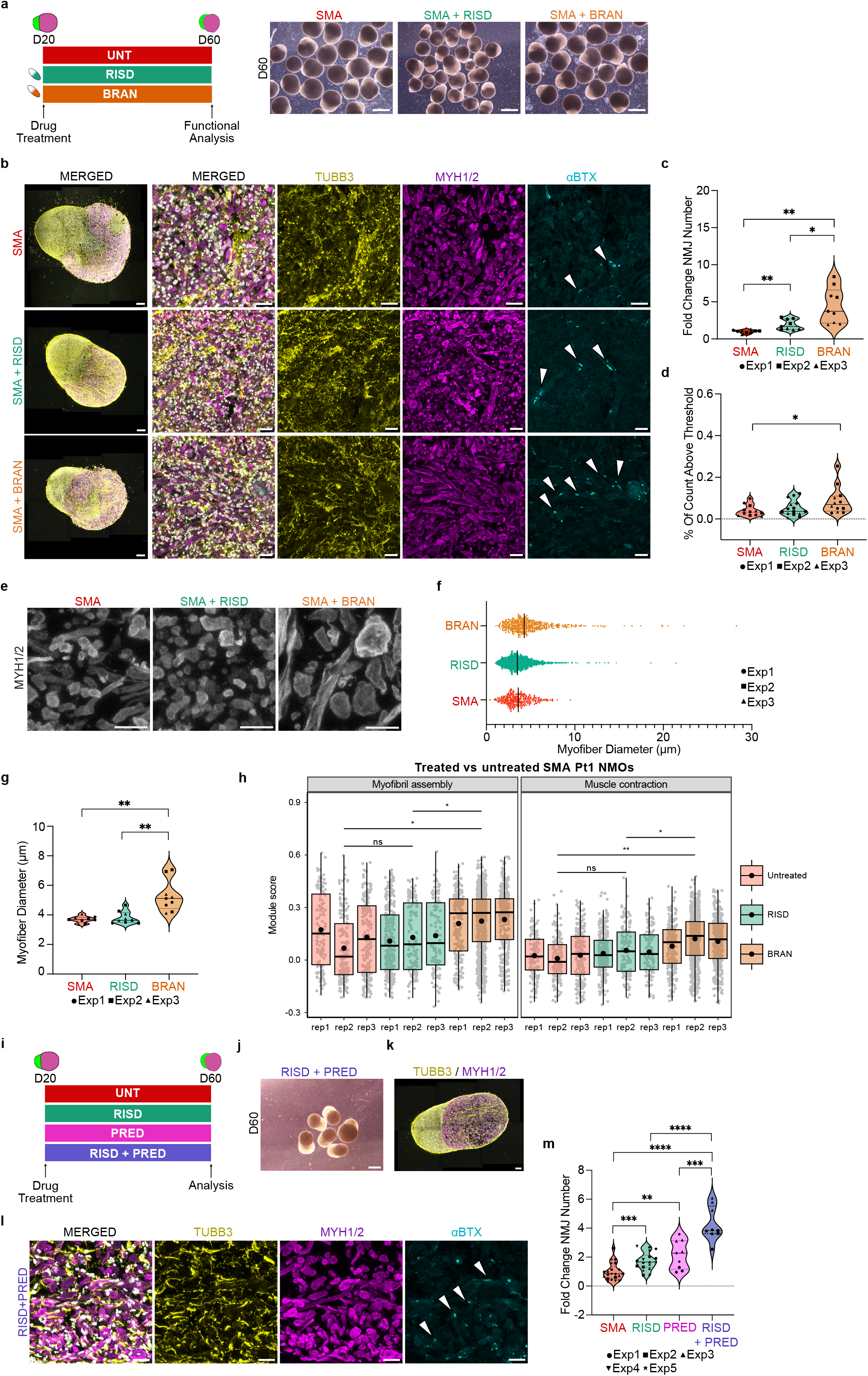
Drug-specific rescue of NMJ formation and myofiber maturation in SMA NMOs. (a) Experimental timeline with representative brightfield images of day 60 SMA Pt1 NMOs (untreated, Risdiplam, Branaplam). (b) Representative immunofluorescence images of day 60 SMA Pt1 NMOs untreated or treated with Risdiplam or Branaplam, stained for neurons (TUBB3, yellow), muscle (MYH1/2, magenta), and acetylcholine receptor clusters (αBTX, cyan). Arrowheads indicate αBTX clusters. (c) Quantification of NMJ number normalised to untreated SMA NMOs. (d) Quantification of NMO contractile function measured as the percentage of counts above threshold, reflecting contraction frequency. (e) Higher magnification images of MYH1/2-stained myofibers across treatment groups. (f) Distribution of myofiber diameters across groups. (g) Quantification of mean myofiber diameter across groups. (h) Functional module expression scoring in myonuclei from drug-treated and untreated NMOs hints at a Branaplam-specific enhancement of muscle functions, namely, myofibril assembly and muscle contraction. Gene module scores were calculated using Seurat v5.3.1 *AddModuleScore* function. (i) Experimental timeline of combined Risdiplam (250nM) and Prednisolone (10μM) treatment. (j) Representative brightfield images of day 60 NMOs treated with Risdiplam and Prednisolone. (k, l) Representative immunofluorescence images of day 60 SMA Pt1 NMOs treated with Risdiplam plus Prednisolone, stained for neurons (TUBB3, yellow), muscle (MYH1/2, magenta), and acetylcholine receptor clusters (αBTX, cyan). Arrowheads indicate αBTX clusters. (m) Fold change in NMJ number relative to untreated SMA NMOs across Risdiplam, Prednisolone and combinatorial Risdiplam plus Prednisolone conditions. Scale bars: (a) 1 mm, (b) 150 µm and 25 µm, (e) 20 µm, (j) 1 mm, (k) 100 µm, (l) 25 µm. Data in panels c, d, g, and m are presented as violin plots displaying the median, quartiles, and individual data points. Data in panel f is shown as a scatter dot plot displaying the mean and individual data points. Each dot represents one NMO (c, d, g, m) or one myofiber (f). Different symbols denote independent experiments, with N = 3 and n = 3 for all conditions in panels c, d, f, and g. For panel m, untreated and Risdiplam groups include N = 7 and n = 3, and Prednisolone and combinatorial Prednisolone + Risdiplam treatments include N = 3 and n = 3. Statistical significance was assessed using two-sided Welch’s t-test (*P < 0.05; **P < 0.01; ***P < 0.001; ****P < 0.0001). For panel h, module scores of 3 biological replicates of untreated (n1=131, n2 = 159, n3 = 207), risdiplam-treated (n1 = 284, n2 = 126, n3 = 107) and Branaplam-treated (n1 = 210, n2 = 774, n3 = 374) nuclei were modeled using a linear mixed-effects model (via lme4 v1.1-37) with treatment group and replicate as fixed effects and sample as a random intercept. Tukey-adjusted pairwise comparisons were subsequently performed using emmeans v2.0.0 (*P < 0.05; **P < 0.01). RISD = Risdiplam, BRAN = Branaplam, PRED = Prednisolone.

To test whether targeting muscle-specific pathways could enhance structural and functional outcomes, we combined Risdiplam with Prednisolone, a corticosteroid known to promote muscle growth^42^. Co-treatment significantly increased both NMJ number and myofiber diameter compared to Risdiplam alone (Fig.5i-m and Extended Data Fig.8g), indicating that enhancing muscle intrinsic programs can potentiate the effects of SMN restoration. Prednisolone alone also improved muscle growth and NMJ number (Fig.5m and Extended Data Fig.8f-h), although the effect on NMJs was more pronounced in combination with Risdiplam, indicating synergistic enhancement of neuromuscular function (Fig.5m and Extended Data Fig.8g,h).

Collectively, these findings demonstrate that SMN2 splicing modulators exert distinct lineage-specific effects despite targeting the same pathway. While both Risdiplam and Branaplam restore SMN levels and increase NMJ formation, their downstream effects differ. In neurons, treatment is associated with partial reversal of disease-related programs, whereas in skeletal muscles, Branaplam induces a broader response that includes potentiation of genes linked to muscle growth and contractile function. These differences are reflected functionally, where improved neuromuscular output is associated with enhanced muscle maturation rather than NMJ number alone. Furthermore, combinatorial treatment with Prednisolone supports the contribution of muscle-intrinsic pathways to functional recovery.

## Discussion

The development of therapies for spinal muscular atrophy (SMA) has been hindered by the lack of human models that capture neuromuscular interactions and patient heterogeneity. Here, we establish patient-derived neuromuscular organoids (NMOs) as a self-organising, multi-lineage system that recapitulates key molecular, structural, and functional hallmarks of SMA. By integrating automated differentiation workflows and cryobanking of neuromesodermal progenitors, this platform enables standardised and scalable generation of NMOs, reduces operator-dependent variability, and allows scalable generation of NMOs from multiple patient lines. Notably, this platform combines automation, cryobanking, and multi-lineage organoid generation in a unified workflow, enabling reproducible and scalable production of functional neuromuscular systems compatible with downstream molecular and functional analyses. This system enables high-resolution, cell-type-resolved analysis of disease mechanisms and therapeutic responses within an integrated neuromuscular context. This work extends current organoid methodologies by introducing an automated and scalable workflow for generating multi-lineage functional neuromuscular organoids.

Our findings support a multi-lineage model of SMA pathology. Although all patient-derived NMOs exhibited reduced NMJ numbers, structural deficits alone did not explain functional impairment. The most severely affected line (Pt1) displayed profound contractile deficits and failed to support myofiber growth, whereas Pt2 and Pt3 maintained functional maturation despite similar NMJ loss. These results indicate that muscle-intrinsic processes, including fibre growth, maturation, and survival, contribute to disease severity, shifting the perspective from a predominantly neuron-centric model toward a coordinated dysfunction across neural and muscle compartments.

Using this platform, we dissected the effects of two clinically relevant SMN2 splicing modulators. While both Risdiplam and Branaplam restored SMN expression and increased NMJ number, their downstream effects diverged across lineages. In neuronal populations, both compounds induced partially overlapping transcriptional changes in synaptic and functional gene programs. In contrast, muscle responses were highly divergent. Branaplam induced broader transcriptional changes, including activation of myogenic and structural programs associated with muscle maturation, whereas Risdiplam showed comparatively limited effects in the muscle compartment. These lineage-dependent differences were associated with functional outcomes, with only Branaplam promoting myofiber growth and improving contractile activity.

These findings indicate that SMN restoration and increased NMJ number alone may not be sufficient to achieve functional recovery. Instead, engagement of muscle-intrinsic maturation programs emerges as a key determinant of improved neuromuscular function. While our data identify a strong association between muscle maturation and functional recovery, direct perturbation experiments will be required to establish causality. In this context, Branaplam, despite its lack of clinical viability, serves as a functional probe to reveal lineage-specific transcriptional programs that are not engaged by SMN restoration alone. These observations are consistent with the variable therapeutic responses observed in patients and highlight a key limitation of strategies that focus exclusively on SMN restoration. Together, these findings provide a framework for evaluating therapeutic responses in human neuromuscular context, with the potential to improve prediction of cell-type-specific drug effects.

To address this limitation, we implemented a combinatorial approach by pairing Risdiplam with Prednisolone, a compound known to promote muscle growth^43^. Co-treatment enhanced NMJ number and increased myofiber diameter compared to Risdiplam alone, demonstrating that targeting muscle-intrinsic pathways can augment the phenotypic effects of SMN restoration. These findings support a model in which effective therapeutic intervention in SMA may benefit from coordinated targeting of both neural and muscle compartments.

Several limitations should be considered. Although NMOs were generated from three SMA patient lines, detailed drug response analyses were primarily performed in the most severely affected line (Pt1). While the use of isogenic controls would further strengthen these findings, precise genome editing of the SMN1/SMN2 locus remains technically challenging due to its repetitive structure. Moreover, although the automated workflow enables scalable organoid generation, downstream analyses remain constrained by organoid size and complexity, limiting high-throughput imaging for large drug screening applications and functional assays. Future efforts to miniaturise organoid formats and integrate advanced imaging and analysis pipelines may further enhance the scalability of this platform.

Collectively, our study establishes SMA NMOs as a mechanistically informative and translationally relevant human platform that bridges disease modelling with therapeutic testing. By resolving lineage-specific disease mechanisms and drug responses within a human multi-lineage system, this approach highlights muscle maturation, rather than NMJ number alone, as a key determinant of functional recovery, with implications for therapeutic efficacy that are not captured in reductionist models. Beyond SMA, this platform provides a framework for modelling complex neuromuscular diseases and for the combinatorial design of multi-lineage therapeutic strategies.

## METHOD DETAILS

### Human pluripotent stem cell lines and culture conditions

The female H9, and the male H1 human embryonic stem cell lines were obtained from WiCell, and approved for use in this project by the Regulatory Authority for the Import and Use of Human Embryonic Stem Cells in the Robert Koch Institute (AZ:3.04.02/0123). The male WTC-TTNGFP iPSC line (UCSFi001-A-27) was obtained from the Allen Institute for Cell Science (AICS). The SMA patient lines included SMA Pt1 (SMAE1C4)^8^, SMA Pt2 (CS84iSMA, Cedars Sinai), and SMA Pt3 (CS86iSMA, Cedars Sinai). All lines were maintained in mTESR1 medium (Stem Cell Technologies) on Geltrex LDEV-Free hESC-Qualified Reduced Growth Factor Basement Membrane Matrix (Life Technologies) at 37 °C, 5%CO_2_. Cell lines were checked for normal karyotype and were mycoplasma-free. The cells were passaged twice a week using Versene solution (Thermo Fisher) or ReLeSR™ (Stem Cell Technologies).

### Generation of NMOs

NMOs from the WTC-TTNGFP (C1), H1 (C2), H9, SMA Pt1 (SMAE1C4) ^8^, SMA Pt2 (CS84iSMA, Cedars Sinai), and SMA Pt3 (CS86iSMA, Cedars Sinai) PSC lines were generated following the protocol described by Martins et al^7^. The quality of the NMOs was assessed at various stages of differentiation. Key quality check points included the elongation of the NMO with initial segregation of mesodermal and neuronal compartments on day 5, and clear compartmentalization with spontaneous contraction observed around day 50 in control lines. Only NMOs that met this quality criteria were selected for further analysis. At least three independent differentiations for each cell line were performed and analyzed.

### Automated generation of NMOs

For large-scale generation of NMOs, a semi-automated workflow was implemented using a customized liquid-handling robot, based on the Biomek i7 platform. The system, termed Fast Reliable Experimental Design (FRED) was equipped with an integrated CO₂ incubator (Cytomat™, Thermo Fisher Scientific), microplate centrifuge (Agilent), and automated cell counter (Vi-CELL™ XR, Beckman Coulter). Custom protocols for neuromesodermal progenitor (NMP) differentiation, NMO formation, and maintenance were developed and executed using the Biomek software environment. For NMP generation, human induced pluripotent stem cells (hiPSCs) cultured on Geltrex™ -coated OneWell plates (Greiner Bio-One) were dissociated into single cells using TrypLE™ (Gibco, Thermo Fisher Scientific) and re-plated at defined densities. Subsequent differentiation steps and media changes were carried out by FRED according to a standardized schedule, optimized for reproducibility and scalability.

### Cryopreservation and thawing of NMPs

At the end of differentiation, NMPs were washed once with 1× PBS and incubated with Accutase™(Sigma-Aldrich) or TrypLE™ (Gibco, Thermo Fisher Scientific) for 5 minutes at 37°C to obtain a single-cell suspension. The enzymatic reaction was stopped by adding DMEM/F-12 (Gibco, Thermo Fisher Scientific), and the cell suspension was transferred to a 15 mL conical tube. Cells were centrifuged at 2000 rpm for 4 min at room temperature, the supernatant was removed, and the pellet was resuspended in 1 mL DMEM/F-12. After counting viable cells using Trypan blue exclusion, on a Countess^TM^ automated cell counter (Thermo Fisher Scientific), cells were resuspended in Bambanker™ cryopreservation medium (Nippon Genetics) and aliquoted for freezing. Cryovials were placed at –80°C overnight before transfer to liquid nitrogen for long-term storage. For thawing, cryovials were briefly incubated in a 37°C water bath until just thawed. The contents were immediately transferred to pre-warmed DMEM/F-12, followed by centrifugation at 2000 rpm for 4 minutes. The supernatant was discarded, and the cells were resuspended in N2B27 medium ^7^. After counting, cells were seeded into 96-well U-bottom ultra-low attachment plates (Nunc, Thermo Fisher Scientific) for NMO generation.

### Reverse transcription - Quantitative PCR Analysis

Total RNA was isolated from NMPs in monolayer using Trizol^TM^ reagent according to the manufacturer’s instructions. First strand cDNA synthesis of 2 µg total RNA was performed with SuperScript™ VILO™ (Thermo Fisher Scientific) and amplified using Platinum SYBR-Green (Thermo Fisher Scientific). For QPCR the Applied Biosystems QuantStudio 7 Pro Real-Time PCR system was used. PCR primers were designed using NCBI Primer-Blast software, using exon-spanning junctions. Expression values for each gene were normalized against GAPDH, and plotted as delta Ct (*ΔCt*).

### Drug treatment of NMOs

NMOs derived from the SMAE1C4, CS84, and CS86 iPSC lines were treated with Branaplam (MCE, HY-19620) or Risdiplam (MCE, HY-109101) starting on day 20.

For the treatment, 30 NMOs were cultured in 6 mL of N2B27 medium supplemented with 40 nM Branaplam or 250 nM Risdiplam. For combinatorial treatment of SMA Pt1 NMOs 10 µM Prednisolone was added to either Risdiplam-treated cultures or cultures without Risdiplam. NMOs that did not receive any drug treatment served as untreated controls. The medium was changed every other day. NMOs were harvested after 10 and 40 days of treatment and subsequently processed for single-nucleus RNA sequencing, immunofluorescence analysis, protein extraction and contraction assays. The functionality of NMJs was tested by pharmacological blocking with Curare as described before^7^.

### Single-nucleus RNA sequencing (snRNA-seq) of NMOs

NMOs derived from the WTC-TTNGFP, H9, SMAE1C4, CS84, and CS86 iPSC lines were collected on day 30 for snRNA-seq. Additionally, NMOs from the SMAE1C4 iPSC lines treated with Branaplam or Risdiplam were collected on day 30 for snRNA-seq. For each genotype or treatment group, 3-5 NMOs were pooled and snap-frozen.

Single nuclei were extracted by mechanical dissociation in NP-40 lysis buffer composed of 10 mM Tris-HCl (pH 7.4), 10 mM NaCl, 2.69 mM MgCl_2_, 0.1% NP-40, 1 mM DTT, 1 mM complete EDTA-free protease inhibitor, and 1U/µL Protector RNase inhibitor. After centrifugation at 500 g for 5 min at 4°C, the pellet was resuspended in wash buffer composed of 1% BSA in 1x PBS supplemented with 0.4 U/µL Takara RNase inhibitor. Isolated nuclei were stained with DAPI, filtered through a 40 µm Flowmi™ (Sigma-Aldrich) cell strainer, and sorted by FACS. Following FACS, 10,000 nuclei were processed for sequencing according to the manufacturer’s instructions using the Chromium Next GEM Single Cell 3ʹ Reagent Kits v3.1 (Dual Index). Samples were sequenced on a NovaSeq X Plus system (Illumina) on 10B lanes, Runmode: 28+10+10+90.

### snRNA-seq data analysis

Sequenced reads were aligned to the GRCh38 reference genome and counted using 10x Genomics Cell Ranger v7.1.0 analysis pipeline. All downstream analyses of the read count matrix were performed using Seurat v5.3.1. Ambient RNA correction of all samples was performed using CellBender v0.3.2. Nuclear fraction scores were computed using DropletQC v0.0.0.9000. Low-quality nuclei with low library size/feature count, low nuclear fraction scores or high mitochondrial read percentages were excluded using a per-sample adaptive thresholding approach. Nuclear doublets/aggregates were identified and excluded using scDblFinder v1.20.2. The filtered count matrices were normalized and variance-stabilized using Seurat sctransform v2. To correct for sample-dependent batch effects, the principal components (PCs) were corrected and integrated using Harmony v1.2.4. Following the Seurat standard workflow, dimensionality reduction and clustering of nuclei were performed using the Leiden modularity optimization algorithm (resolution = 0.4). The resulting 18 clusters were cell-type annotated by mapping to published single-cell atlases of human developing and adult spinal cord and adult skeletal muscle tissues. Cluster biomarkers were identified for cell identity validation by Wilcoxon testing of gene overexpression in one cluster versus all others. Differential gene expression analyses between SMA and control and between drug-treated and untreated SMA Pt1 NMOs were performed in various cell clusters via a pseudobulk approach using DESeq2 v1.46.0. Genes with an absolute fold change > 1.5 following fold change shrinkage via apeglm v1.28.0 and a Benjamini-Hochberg (BH) adjusted P-value < 0.05 were declared significantly differentially expressed (Supplementary Table 1). Gene set enrichment analysis was performed using fgsea v1.32.4 and enrichR v3.4 (Supplementary Table 2). Module scores for gene sets of interest were calculated using Seurat *AddModuleScore* function. Differences in scores between treatment groups were statistically tested using linear mixed-effects models (treatment group and replicate as fixed-effects and sample as a random intercept) implemented with lme4 v1.1-37 and Tukey-adjusted pairwise comparisons were performed using emmeans v2.0.0. Drug transcriptional reversal effects were assessed by testing the negative and positive enrichment of pre-ranked drug DEG lists for the top 200 SMA-upregulated and downregulated genes, respectively. All analysis code and code used to generate the figures will be made available on GitHub upon publication. All software tools used in the analysis will be made available upon publication.

### Western Blot analysis

Whole protein extraction was carried out using RIPA Lysis Buffer System™ (Santa Cruz). Protein concentrations were determined using the Pierce™ BCA Protein Assay Kit (Thermo Fisher Scientific), following the manufacturer’s instructions in a 96-well format. For SDS-PAGE, 20 µg of protein were loaded onto a 10% SDS-PAGE gel and separated by electrophoresis at 120 V for 1 hour. Proteins were then transferred to PVDF membranes using the Trans-Blot Turbo Transfer System (BioRad) for 10 minutes at 25 V. The membranes were blocked with 3% skimmed milk in TBS with 0.1% Tween 20 (TBST) for 1 hour at room temperature. Following blocking, the membranes were incubated overnight at 4°C with continuous shaking using primary antibodies against SMN (BD Transduction) or GAPDH (Sigma Aldrich). After incubation, membranes were washed in TBST for 1 hour at room temperature, and subsequently incubated with either rabbit-HRP or mouse-HRP secondary antibodies in 3% skimmed milk in TBS for 1 hour at room temperature. Protein detection was performed using the Amersham ECL reagent, with imaging carried out using the ChemiDoc™ MP Imaging System. Band intensities were quantified in Fiji Image J 1.54p (ImageJ) using the gel analysis tool: rectangular ROIs were drawn around each lane, and the area under each band peak was measured. SMN band intensities were normalized to the corresponding GAPDH signal from the same lane. Final SMN expression values were expressed relative to untreated SMA control samples.

### NMO size measurement

On day 1 and day 5 of NMO development, size measurements were performed using the Opera Phenix™ Plus High Content Screening System (PerkinElmer) in brightfield mode with a 5x objective. Images were acquired and analyzed using Harmony5.2 software (PerkinElmer), which provided measurements of NMO area and elongation. NMO size distribution was calculated by normalizing individual NMO sizes to the mean NMO size within each differentiation batch. NMO structures with values below 0.5 were excluded from the analysis. For size measurements on days 10, 20, and 30, NMOs were imaged using an Olympus SZX16 stereomicroscope at 0.7x magnification. Size analysis was conducted in Fiji Image J 1.54p (ImageJ) by manually delineating each organoid using the freehand selection tool, and the measured area (in pixels) was recorded via the ROI Manager. Fused or unsegregated NMOs were excluded from the analysis.

### Immunofluorescence of NMOs

Immunofluorescence staining of NMOs was performed following the previously described protocol ^7^. Organoids were fixed, permeabilised, and blocked as previously described. Primary antibodies used in this study will be made available upon publication, including their respective host species, dilutions, and suppliers. Detection of primary antibodies was achieved using species-specific secondary antibodies conjugated to Alexa Fluor 488, 568, or 647 (Invitrogen, Thermo Fisher Scientific). Nuclei were counterstained with DAPI (Sigma-Aldrich).

### Image analysis

NMJ number and size, myofiber count and diameter, and NMJ innervation were assessed from confocal images acquired using a LEICA SP8 microscope with a 63x oil immersion objective. NMJs were identified via αBTX^+^ staining and quantified either manually in Fiji or using a semi-automated pipeline based on ilastik segmentation and Python analysis (ilastik 1.4.0.post1, Python 3.9.21). To quantify neural and muscle tissue proportions, NMO size was measured in Fiji by calculating the DAPI-, TUBB3-, and MSF-positive areas as previously described^7^. NMJ size was defined as the major axis length of each αBTX-labeled region, and innervation was determined by overlap with TUBB3 staining. Myofibers were counted manually from MSF-stained images using Fiji’s Multi-point tool and NMJ number was normalised to myofiber count per field of view. Myofiber diameter was measured as the minimum Feret’s diameter from cross-sectional MSF-stained images using straight-line ROI measurements in Fiji (Image J 1.53q or 1.54p). Quantification of NMJ number, size, myofiber count, and NMJ innervation was performed using three fields of view per NMO. Fold change in myofiber growth was calculated by dividing the average myofiber size at each time point (day 30 and day 60) by the average myofiber size at day 30 for each cell line.

For semi-automated quantification of NMJs and their innervation, a custom image analysis pipeline was developed using ilastik and Python (ilastik 1.4.0.post1, Python 3.9.21). Confocal images stained for αBTX and TUBB3 were first processed using ilastik, to generate pixel classification models and corresponding masks, which were exported as NumPy arrays (*.npy). In the Python-based analysis, αBTX-positive regions were labelled, and NMJ size was calculated as the major axis length of an ellipse fitted to each labelled region using the scikit-image library. For innervation analysis, TUBB3 masks were dilated by 2 pixels and overlaid with αBTX masks to identify regions of signal overlap. Only NMJs larger than 2 µm in length were considered for final quantification. Output data and validation images were automatically exported for downstream analysis. Fold change in NMJ size was calculated by dividing the average NMJ size at each time point (day 30 and day 60) by the average NMJ size at day 30 for each cell line.

Apoptosis in NMO sections was quantified in Fiji (Image J 1.53q) using cleaved CASP3 immunostaining. Regions of interest (ROIs) were defined for the whole organoid (DAPI), muscle (Desmin), motor neurons (SMI32), and neural tissue (DAPI excluding Desmin). Each channel was converted to 8-bit, filtered (Gaussian blur or min/max), and auto-thresholded to generate ROIs. Cleaved CASP3-positive areas were segmented using ilastik as described above. Apoptosis in specific regions was determined by intersecting cleaved CASP3 ROIs with muscle or neural ROIs (AND operation) and area and intensity values were extracted for analysis.

### Functional characterization of NMOs

Spontaneous contraction of NMOs was evaluated based on live recordings, using a confocal microscope (Leica SP8), in incubation mode (37 °C and 5% CO2), and a semi-automated analysis pipeline we have previously developed^44^. Briefly, on the day of the assay, NMOs (day 54-65) from each condition (WTC-TTNGFP), CS84, CS86, or SMAE1C4 NMOs +/- drug) were collected in dedicated wells of a 24-well plate (one organoid per well) with fresh N2B27 media.

For live imaging, most of the media was removed from the well (∼20μL of media left for hydration) in order to immobilise the organoid. A brightfield image (10x objective) was acquired of the whole NMO, serving as a map for determining the location of the recordings. In this study, we acquired recordings from three NMO positions: two at the outer edge of the organoids at the interface of the neural and muscular compartments and one at the pole of the muscular compartment (see Fig. 2f). To this end, using 4.5x magnification, a close-up frame was set so that half of it was filled by the organoid and the other half by the background. Then, one to three 10-minute videos of each position of each NMO were acquired. If NMOs exhibited minimal activity (i.e., no or very subtle movements in two-three consecutive 10-min videos), then only two to three recordings were acquired. Live recordings from at least four NMOs from each condition, from each independent differentiation and each condition were acquired.

The above dataset was then processed with a two-stage semi-automated analysis pipeline, as described here^44^. During the first stage, contraction recordings are processed so that changes in the border of the organoid over time are extracted as signals (*.npy files) and visualised as *Time Series* plots. In the second stage, time series signals undergo a pre-processing step, followed by extraction of contraction features using Python packages *ts-fresh.* More specifically, the following features were extracted:

#### Contraction Power

This refers to the absolute energy of our *Time series* signal, *y(t)* (i.e., the displacement in y-axis in *Time series* plots of the NMO border over time). To calculate this, the absolute energy of the signal from the Python package ts-fresh was used, as is commonly used as a proxy for the overall magnitude of a signal across its duration. This refers to the sum of the squared values of a signal, according to the following equation:

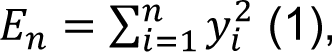

defining *E_n_* as the area under the curve of the squared time series. To account for time resolution (i.e., duration of signal/contraction recording), the above equation was modified by dividing the absolute energy by the duration of the signal (i.e., the number of bins*, n,* multiplied by the time resolution for a discrete signal*, Δt, (empirically set to Δt=0.096s* in this study, corresponding to the sampling interval of the time series). This is then defined as *Contraction power* (μm^2^ s^-1^) and is computed according to the following equation:

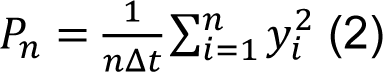

As such, contraction power represents the average value of the squared signal over time, meaning the average intensity of the signal and, thereby, it acts an indirect measure of the NMO contraction strength. *% of count above threshold*: This feature describes the percentage of values in the time series (*y(t)*) that are higher than the given threshold (*th*) below which movements are considered noise. Empirically, in this study, this threshold is set at 0.5 μm. The above features for each contraction video are then extracted as raw data in *.xlsx files for downstream analysis.

### Statistical analysis

All data analysis and visualisation were performed using GraphPad Prism 10. Results are presented either as scatter dot plots showing the mean ± standard deviation and individual data points, or as violin plots displaying the median, quartiles, and individual values. Statistical comparisons were performed using unpaired *t*-tests with Welch’s correction to account for unequal variances. For drug treatment experiments, statistical analyses were conducted by comparing treatment conditions at the same time point within each experimental batch. Quantitative features were extracted from each contraction video and compiled in Excel files as raw data. Statistical significance was tested using an unpaired t-test with Welch’s correction (*\* p < 0.05, ** p < 0.01, *** p < 0.001, **** p < 0.0001*).

### Data availability

All analysis code and code used to generate the figures will be made available on GitHub upon publication.

## Acknowledgements

The authors thank the genomics technology platform at the Max-Delbrück-Center for Molecular Medicine (MDC), Berlin, Germany, for substantial technical support and assistance in this work. We thank Andrea Grybowski, Nicole Grieger, and Michelle Müller for technical support. For the critical reading of the manuscript, we would like to thank Anthony Gavalas. The work was supported by the Max Delbrück Center (MDC), which receives its core funding from the Helmholtz Association. M.G. is funded by the European Research Council (ERC) under the European Union’s Horizon 2020 research and innovation program (GPS-organoids; 101002689), under the European Union’s Horizon 2022 research innovation program (MiniOrgans; 101113481), the Einstein Stiftung Berlin (Einstein Center 3 R, EZ-2020-597-2), the Deutsche Forschungsgemeinschaft (DFG) (400728090; GO 3432/1-2) and the European Molecular Biology Organization Young Investigator program award. L.V.N.N. was supported by an ECRT PhD fellowship. C.M.M. was supported by Marie Skłodowska-Curie grant *e-NeuroMus* (Grant Agreement ID 101149182).

## Author contributions

M.G. conceived the study, including the development of automated NMO generation strategies, designed the experiments, supervised the study, and wrote the manuscript. I.L. designed and performed experiments, established automated NMO generation workflows, conducted data analysis, and prepared the figures. A.G.P. established the generation of SMA NMOs from all patient lines, designed and conducted experiments, and performed data analysis. I.A.E. analysed the snRNA-seq data and prepared the corresponding figures. I.A.M. performed manual and automated NMO generation experiments, drug testing experiments, performed data analysis, and established the pipelines for automated image analysis. L.V.N.N. established the drug testing experiment in SMA NMOs, designed and conducted experiments, and performed data analysis. C.M.M. performed contraction recordings and subsequent analysis. N.F. carried out manual and automated NMO differentiation experiments and provided technical support. I.M.R. performed programming of the liquid handling robot in collaboration with I.L. S.D. provided the automation platform, technical support, and banking of the iPSC lines. C.B., and D.C. developed the code and tool for the contraction analysis under the supervision of M.P. G.J.B. and W.R. provided the SMA patient 1 iPSC line.

## REFERENCES

1 Kim, J., Koo, B. K. & Knoblich, J. A. Human organoids: model systems for human biology and medicine. Nat Rev Mol Cell Biol 21, 571–584 (2020). 10.1038/s41580-020-0259-3

2 Sharma, A., Sances, S., Workman, M. J. & Svendsen, C. N. Multi-lineage Human iPSC-Derived Platforms for Disease Modeling and Drug Discovery. Cell Stem Cell 26, 309–329 (2020). 10.1016/j.stem.2020.02.011

3 Chen, Y. et al. Bridging single cells to organs: Mesoscale modules as fundamental units of tissue function. Cell 188, 6393–6410 (2025). 10.1016/j.cell.2025.10.012

4 He, Z. et al. An integrated transcriptomic cell atlas of human neural organoids. Nature 635, 690–698 (2024). 10.1038/s41586-024-08172-8

5 Gouti, M. et al. In vitro generation of neuromesodermal progenitors reveals distinct roles for wnt signalling in the specification of spinal cord and paraxial mesoderm identity. PLoS Biol 12, e1001937 (2014). 10.1371/journal.pbio.1001937

6 Gouti, M. et al. A Gene Regulatory Network Balances Neural and Mesoderm Specification during Vertebrate Trunk Development. Dev Cell 41, 243–261 e247 (2017). 10.1016/j.devcel.2017.04.002

7 Faustino Martins, J. M., et al. Self-Organizing 3D Human Trunk Neuromuscular Organoids. Cell Stem Cell 26, 172–186 e176 (2020). 10.1016/j.stem.2019.12.007

8 Urzi, A. et al. Efficient generation of a self-organizing neuromuscular junction model from human pluripotent stem cells. Nat Commun 14, 8043 (2023). 10.1038/s41467-023-43781-3

9 Nebol, A. & Gouti, M. A new era in neuromuscular junction research: current advances in self-organized and assembled in vitro models. Curr Opin Genet Dev 87, 102229 (2024). 10.1016/j.gde.2024.102229

10 Henrique, D., Abranches, E., Verrier, L. & Storey, K. G. Neuromesodermal progenitors and the making of the spinal cord. Development 142, 2864–2875 (2015). 10.1242/dev.119768

11 Lunn, M. R. & Wang, C. H. Spinal muscular atrophy. Lancet 371, 2120–2133 (2008). 10.1016/S0140-6736(08)60921-6

12 Sugarman, E. A. et al. Pan-ethnic carrier screening and prenatal diagnosis for spinal muscular atrophy: clinical laboratory analysis of >72,400 specimens. Eur J Hum Genet 20, 27–32 (2012). 10.1038/ejhg.2011.134

13 Lefebvre, S. et al. Identification and characterization of a spinal muscular atrophy-determining gene. Cell 80, 155–165 (1995). 10.1016/0092-8674(95)90460-3

14 Ebert, A. D. et al. Induced pluripotent stem cells from a spinal muscular atrophy patient. Nature 457, 277–280 (2009). 10.1038/nature07677

15 Jha, N. N., Kim, J. K., Her, Y. R. & Monani, U. R. Muscle: an independent contributor to the neuromuscular spinal muscular atrophy disease phenotype. JCI Insight 8 (2023). 10.1172/jci.insight.171878

16 Lorson, C. L., Hahnen, E., Androphy, E. J. & Wirth, B. A single nucleotide in the SMN gene regulates splicing and is responsible for spinal muscular atrophy. Proc Natl Acad Sci U S A 96, 6307–6311 (1999). 10.1073/pnas.96.11.6307

17 Gavrilov, D. K., Shi, X., Das, K., Gilliam, T. C. & Wang, C. H. Differential SMN2 expression associated with SMA severity. Nat Genet 20, 230–231 (1998). 10.1038/3030

18 Finkel, R. S. et al. Nusinersen versus Sham Control in Infantile-Onset Spinal Muscular Atrophy. N Engl J Med 377, 1723–1732 (2017). 10.1056/NEJMoa1702752

19 Pechmann, A. et al. Evaluation of Children with SMA Type 1 Under Treatment with Nusinersen within the Expanded Access Program in Germany. J Neuromuscul Dis 5, 135–143 (2018). 10.3233/JND-180315

20 Arnold, A. S. et al. Reduced expression of nicotinic AChRs in myotubes from spinal muscular atrophy I patients. Lab Invest 84, 1271–1278 (2004). 10.1038/labinvest.3700163

21 Donlin-Asp, P. G. et al. The Survival of Motor Neuron Protein Acts as a Molecular Chaperone for mRNP Assembly. Cell Rep 18, 1660–1673 (2017). 10.1016/j.celrep.2017.01.059

22 Anderson, K. N., Baban, D., Oliver, P. L., Potter, A. & Davies, K. E. Expression profiling in spinal muscular atrophy reveals an RNA binding protein deficit. Neuromuscul Disord 14, 711–722 (2004). 10.1016/j.nmd.2004.08.009

23 Miller, N., Shi, H., Zelikovich, A. S. & Ma, Y. C. Motor neuron mitochondrial dysfunction in spinal muscular atrophy. Hum Mol Genet 25, 3395–3406 (2016). 10.1093/hmg/ddw262

24 Mendell, J. R. et al. Single-Dose Gene-Replacement Therapy for Spinal Muscular Atrophy. N Engl J Med 377, 1713–1722 (2017). 10.1056/NEJMoa1706198

25 Baranello, G. et al. Risdiplam in Type 1 Spinal Muscular Atrophy. N Engl J Med 384, 915–923 (2021). 10.1056/NEJMoa2009965

26 Ratni, H. et al. Discovery of Risdiplam, a Selective Survival of Motor Neuron-2 ( SMN2) Gene Splicing Modifier for the Treatment of Spinal Muscular Atrophy (SMA). J Med Chem 61, 6501–6517 (2018). 10.1021/acs.jmedchem.8b00741

27 Cheung, A. K. et al. Discovery of Small Molecule Splicing Modulators of Survival Motor Neuron-2 (SMN2) for the Treatment of Spinal Muscular Atrophy (SMA). J Med Chem 61, 11021–11036 (2018). 10.1021/acs.jmedchem.8b01291

28 Estevez-Fraga, C., Tabrizi, S. J. & Wild, E. J. Huntington’s Disease Clinical Trials Corner: March 2024. J Huntingtons Dis 13, 1–14 (2024). 10.3233/JHD-240017

29 Ottesen, E. W. et al. Diverse targets of SMN2-directed splicing-modulating small molecule therapeutics for spinal muscular atrophy. Nucleic Acids Res 51, 5948–5980 (2023). 10.1093/nar/gkad259

30 Krach, F. et al. RNA splicing modulator for Huntington’s disease treatment induces peripheral neuropathy. iScience 28, 112380 (2025). 10.1016/j.isci.2025.112380

31 Grass, T. et al. Isogenic patient-derived organoids reveal early neurodevelopmental defects in spinal muscular atrophy initiation. Cell Rep Med 5, 101659 (2024). 10.1016/j.xcrm.2024.101659

32 Quadrato, G. et al. Cell diversity and network dynamics in photosensitive human brain organoids. Nature 545, 48–53 (2017). 10.1038/nature22047

33 Habib, N. et al. Massively parallel single-nucleus RNA-seq with DroNc-seq. Nat Methods 14, 955–958 (2017). 10.1038/nmeth.4407

34 Zheng, G. X. et al. Massively parallel digital transcriptional profiling of single cells. Nat Commun 8, 14049 (2017). 10.1038/ncomms14049

35 Sanes, J. R. & Lichtman, J. W. Induction, assembly, maturation and maintenance of a postsynaptic apparatus. Nat Rev Neurosci 2, 791–805 (2001). 10.1038/35097557

36 Wang, H. S. et al. KCNQ2 and KCNQ3 potassium channel subunits: molecular correlates of the M-channel. Science 282, 1890–1893 (1998). 10.1126/science.282.5395.1890

37 Dickson, B. J. Molecular mechanisms of axon guidance. Science 298, 1959–1964 (2002). 10.1126/science.1072165

38 Kennedy, T. E., Serafini, T., de la Torre, J. R. & Tessier-Lavigne, M. Netrins are diffusible chemotropic factors for commissural axons in the embryonic spinal cord. Cell 78, 425–435 (1994). 10.1016/0092-8674(94)90421-9

39 Bansal, D. et al. Defective membrane repair in dysferlin-deficient muscular dystrophy. Nature 423, 168–172 (2003). 10.1038/nature01573

40 Chal, J. & Pourquie, O. Making muscle: skeletal myogenesis in vivo and in vitro. Development 144, 2104–2122 (2017). 10.1242/dev.151035

41 Charge, S. B. & Rudnicki, M. A. Cellular and molecular regulation of muscle regeneration. Physiol Rev 84, 209–238 (2004). 10.1152/physrev.00019.2003

42 Al Tanoury, Z. et al. Prednisolone rescues Duchenne muscular dystrophy phenotypes in human pluripotent stem cell-derived skeletal muscle in vitro. Proc Natl Acad Sci U S A 118 (2021). 10.1073/pnas.2022960118

43 Baltgalvis, K. A., Call, J. A., Nikas, J. B. & Lowe, D. A. Effects of prednisolone on skeletal muscle contractility in mdx mice. Muscle Nerve 40, 443–454 (2009). 10.1002/mus.21327

44 Moysidou, C.-M. et al. Guided maturation of human neuromuscular organoids via electrical stimulation. bioRxiv, 2025.2010.2030.685366 (2025). 10.1101/2025.10.30.685366

