## Supporting Information for "A scalable human neuromuscular organoid platform enables lineage-specific analysis of drug responses in spinal muscular atrophy"

**Extended Data Figure 1. Automated workflows support scalable generation of NMOs.**

(a) Brightfield images of day 1 NMOs generated by automated and manual workflows in a 96-well format.

(b) Quantification of NMO size on day 1 generated by automated and manual workflows in a 96-well format. Three independent users performed the experiments. Each data point represents one NMO.

(c) Brightfield images of day 5 NMOs generated by automated and manual workflows in a 96-well format.

(d) Quantification of NMO size on day 5 generated by automated and manual workflows in a 96-well format. Three independent users performed the experiments. Each data point represents one NMO.

(e, f) Quantification of day 5 organoid size uniformity (e) and length-to-width ratios (f), demonstrating comparable morphology across conditions. M = manual, F = frozen and A = automated.

(g) Representative brightfield images of day 60 control NMOs derived from frozen NMPs.

(h) Representative immunofluorescence images of day 60 control NMOs derived from frozen NMPs stained for neurons (TUBB3, yellow), muscle (MYH1/2, magenta), and acetylcholine receptor clusters (αBTX, cyan). Arrowheads indicate αBTX clusters.

Scale bars: (g) 1 mm, (h) 250 µm and 25 µm.

Data in b, d are presented as scatter dot plots showing the mean ± s.d. along with individual data points. Each dot represents one NMO. Different symbols denote independent experiments performed by different users. Data in panels e and f are presented as violin plots displaying median, quartiles, and individual data points. Different symbols indicate independent experiments with N=3 for all conditions in panels e and f. Each dot represents one NMO.

**Extended Data Figure 2. Early morphological characterization of SMA neuromuscular organoids (NMOs).**

(a) Overview table of the six PSC lines used in the study including clinical characteristics.

(b) Brightfield images of day 5 SMA NMOs seeded in a 96-well format. Insets: representative day 5 NMOs from SMA patient lines.

(c, d) Quantification of NMO size distribution (c) and elongation (d) on day 5. Each dot represents one NMO.

Data in panels c and d are presented as scatter dot plots displaying the mean ± s.d. along with individual data points.

**Extended Data Figure 3. Cellular composition and SMN expression in early SMA NMOs.**

(a) Representative immunofluorescence images of day 20 SMA NMOs stained for muscle markers (DESMIN, MYOD1, PAX7) and neuronal markers (TUBB3, ISL1, SOX1). Nuclei are counterstained with DAPI.

(b) Western blot and quantification of SMN protein (34 kDa) expression in day 30 NMOs from control (C1) and SMA patient lines (Pt1–Pt3). GAPDH (36 kDa)was used as a loading control.

(c) Representative immunofluorescence images of day 50 NMOs stained for neurons (TUBB3, green) and muscle (MYH1/2, magenta), with merged images shown below.

(d,e) Percentage of neural and muscle tissue in NMOs.

Scale bars: (a) 10 µm, (c) 100 µm.

Data in panels d and e are presented as violin plots displaying the median, quartiles, and individual data points. Different symbols represent independent experiments, with N = 3 and n = 3 for all conditions.

**Extended Data Figure 4. Quantitative analysis of NMJ maturation, myofiber development and contractile function in control and SMA neuromuscular organoids (NMOs).**

(a, b) Quantification of NMJ size at day 30 (a) and day 60 (b) in control (C1, C2) and SMA NMOs (Pt1–Pt3).

(c) Fold change in NMJ size between day 30 and day 60.

(d) Ratio of innervated NMJs at day 60, defined as the proportion of $aBTX^+$ clusters overlapping with TUBB3 neuronal projections relative to the total number of $aBTX^+$ clusters.

(e) (Quantification of NMJ number ($\alpha BTX^+$ clusters) per 63× field of view in control and SMA NMOs generated using manual and automated workflow.

(f, g) Quantification of myofiber diameter at day 30 (f) and day 60 (g).

(h) Distribution of myofiber diameters per line at day 60. Each datapoint represents one myofiber.

(i) Quantification of myofiber number per FOV at day 60 in control (C1, C2) and SMA NMOs (Pt1–Pt3).

(j) Quantification of NMO contractile activity expressed as Contraction power ($\mu m^2/s$).

Data in panels a–g and i,j are shown as violin plots displaying the median, quartiles, and individual data points. Data in panel h is presented as a scatter dot plot showing the mean and individual data points. Different symbols indicate independent experiments, with N = 3 and n = 3 for all conditions in panels a - j. Each data point represents one NMO (a-g, i,j) or one myofiber (h). Statistical significance was assessed using Welch's t-test (*P < 0.05; **P < 0.01; ***P < 0.001; ****P < 0.0001).

**Extended Data Figure 5. Increased apoptosis in muscle regions of SMA neuromuscular organoids (NMOs).**

(a) Representative immunofluorescence images of day 50 NMOs stained for neurons (TUBB3, green), muscle (ACTN2, magenta), and apoptotic cells (cleaved Caspase-3, yellow). Arrowheads indicate Cleaved Caspase-3 positive cells.

(b) Quantification of apoptotic area in the muscle region at day 50.

(c) Representative immunofluorescence images of day 50 NMOs stained for neurons (Neurofilament-H, green), motor neurons (ChAT, turquoise), and apoptotic cells (Cleaved Caspase-3, yellow). Arrowheads indicate cleaved Caspase-3–positive cells.

(d) Quantification of apoptotic area in the neuronal region at day 50.

Scale bars: (a, c) 100 µm and 10 µm.

Data in panels b and d are shown as violin plots displaying the median, quartiles, and individual data points. Different symbols indicate independent experiments, with N = 3 and n = 3 for all conditions in panels b and d. Each data point represents one NMO. Statistical significance was assessed using two-sided Welch's t-tests (*P < 0.05; **P < 0.01; ***P < 0.001; ****P < 0.0001).

**Extended Data Figure 6. Characterisation of nuclei clusters and cell type composition of day 30 control and SMA NMOs.**

(a) UMAP representations of the two control NMO lines combined (n = 12,365) and the three SMA lines: SMA Pt1 (n = 3,592), SMA Pt2 (n = 3,085) and SMA Pt3 (n = 2,949),showing comparable cell-type composition across control and SMA NMOs.

(b) UMAPs revealing expression landscapes of key cell type markers in NMOs, namely PAX6 (neural progenitors), ELAVL4 (neurons), GFAP (astroglia), PAX7 (satellite cells), ACTN2 (skeletal muscles) and PDGFRA (fibroblasts).

(c) Expression of key cell-type markers across neuroglial, muscle and fibroblast clusters. These include markers of cell proliferation (PCNA, TOP2A, MCM6 & MKI67), neural progenitors (PAX6, HES5 & FABP7), immature neurons (HES6 & DLL3), mature neurons (ELAVL4, ELAVL3, STMN2, NSG2, RBFOX3 and MAP2), excitatory neurons (SLC17A6), inhibitory neurons (GAD1 & GAD2), astroglia (GFAP, DKK1, EFNB3 and SCG2), Schwann cells (CDH19, ASPA and MPZ), satellite cells (PAX7, PITX2 and SYTL2), myocytes (MYMX, SGCA, MYOD1, MYOG, CDH15 and CHRNB1), muscle fibers (CHRND, TTN, MYBPH, ACTA1, ACTN2 and UNC45B) and fibroblasts (FAP, PDGFRA, COL1A1, COL1A2 &

COL3A1). Dot colour and size represent the average marker expression level and percentage of marker-expressing nuclei per cluster, respectively.

**Extended Data Figure 7. Transcriptional effects of Risdiplam and Branaplam in day 30 NMOs.**

(a) Experimental timeline for treatment of SMA patient-derived NMOs (Pt2, Pt3) with Risdiplam, Branaplam, or no treatment from day 20 to day 30 of differentiation.

(b) Western blot of SMN protein (34 kDa) levels in treated and untreated SMA Pt2 and Pt3 NMOs. GAPDH (36 kDa) was used as a loading control. A control line was included for reference.

(c) Quantification of the SMN protein levels following treatment with Risdiplam and Branaplam relative to untreated conditions.

(d) Neuroglial and mesodermal cell type compositions of drug-treated and untreated NMO lines, presented as the percentage of each cell type withing each lineage. An increase in the proportion of skeletal muscle cells is observed following Branaplam treatment.

(e) Numbers of genes whose expression is significantly reversed or potentiated by each drug across major cell types. Overall, drug treatment was associated with a higher number of reversed than potentiated genes, except for the effect of Branaplam on skeletal muscles.

(f) Consistent with e, gene set enrichment analysis shows that Risdiplam and Branaplam treatment is generally associated with negative enrichment of the top 200 SMA-upregulated genes and positive enrichment of the top 200 SMA-downregulated genes, consistent with reversal of disease-associated signatures across cell types. In contrast, skeletal muscle following Branaplam treatment shows enrichment patterns consistent with potentiation rather than reversal. The significance cutoff is set to a Benjamini-Hochberg-adjusted P-value $< 0.05$.

(g) Representative examples of drug-reversed and potentiated genes in neurons and skeletal muscle are displayed together with their associated cellular functions. Genes are annotated as reversed or potentiated based on the direction of expression change following drug treatment in relation to the direction of change in SMA versus control. If gene expression follows in the same direction in both comparisons, it is labelled as “potentiated”, otherwise, it is labelled as “reversed”. Dot color indicates $\log_2$ fold change (treated versus untreated), and dot size represents the percentage of gene-expressing nuclei in each treatment group.

(h) A representation of Risdiplam and Branaplam off-targets that were differentially expressed following drug treatment across the different cell types. These are genes that were previously reported to undergo aberrant splicing and showed expression changes following drug treatment (Ottesen et al., 2023), consistent with observations in the present study.

Data in panel c are presented as violin plots displaying the median, quartiles, and individual data points. Different symbols indicate different SMA patient lines. Statistical significance was assessed using Welch's t-test (*P < 0.05; **P < 0.01).
RISD = Risdiplam, BRAN = Branaplam.

**Extended Data Figure 8. Drug treatment differentially modulates NMJ structure and contractile function in SMA NMOs.**
(a, b) Quantification of NMJ size in untreated, Risdiplam-treated, and Branaplam-treated SMA NMOs at day 60.
(c) Representative contraction time-series plots of SMA NMOs following drug treatment.
(d) Quantification of contractile activity expressed as *Contraction power* ($\mu m^2/s$ in Risdiplam- or Branaplam-treated NMOs.
(e) Experimental timeline of Prednisolone-only treatment from day 20 to day 60 with representative brightfield images of day 60 SMA Pt1 NMOs untreated or Prednisolone-treated.
(f) Representative immunofluorescence images of day 60 SMA Pt1 NMOs untreated or Prednisolone-treated NMOs, stained for neurons (TUBB3, green), muscle (MYH1/2, magenta), and acetylcholine receptor clusters (αBTX, cyan). Arrowheads indicate αBTX clusters.
(g, h) Quantification of fold change in myofiber diameter (g) and NMJ size (h) in untreated, Risdiplam, Prednisolone and combinatorial Risdiplam plus Prednisolone conditions.
Scale bars: (e) 1 mm, (f) 25 µm.
Data in panels a, d, g, and h are shown as violin plots showing median, quartiles, and individual NMO data points. Each data point represents one NMO. Different symbols indicate independent experiments. For panels a and d, all conditions include N=3, n=3. For panels g and h, the untreated and Risdiplam groups include N=7, n=3, whereas the Prednisolone and Risdiplam plus Prednisolone groups include N=3 and n=3. Statistical significance was determined using two-sided Welch's t-test; *P < 0.05, **P < 0.01, ***P < 0.001, ****P < 0.0001.
RISD = Risdiplam, BRAN = Branaplam, PRED = Prednisolone.

**Supplementary Tables**

Supplementary Table 1: DEG lists with significant genes identified across all comparisons

Supplementary Table 2: GSEA lists with significant gene sets for all comparisons

**Supplementary Movies**

Movie S1: Representative video of control NMO spontaneous contraction.

Movie S2: Representative video of SMA Pt1 NMO spontaneous contraction.

Movie S3: Representative video of SMA Pt2 NMO spontaneous contraction.

Movie S4: Representative video of SMA Pt3 NMO spontaneous contraction.

Movie S5: Representative video of SMA Pt2 NMO spontaneous contraction, not treated with curare.

Movie S6: Representative video of SMA Pt2 NMO spontaneous contraction during curare treatment.

Movie S7: Representative video of SMA Pt3 NMO spontaneous contraction, not treated with curare.

Movie S8: Representative video of SMA Pt3 NMO spontaneous contraction during curare treatment.

Movie S9: Representative video of untreated SMA Pt1 NMO spontaneous contraction.

Movie S10: Representative video of Risdiplam-treated SMA Pt1 NMO spontaneous contraction.

Movie S11: Representative video of Branaplam-treated SMA Pt1 NMO spontaneous contraction.
