## Extended Data Figures for "A scalable human neuromuscular organoid platform enables lineage-specific analysis of drug responses in spinal muscular atrophy"

#### Extended Data Figure 1

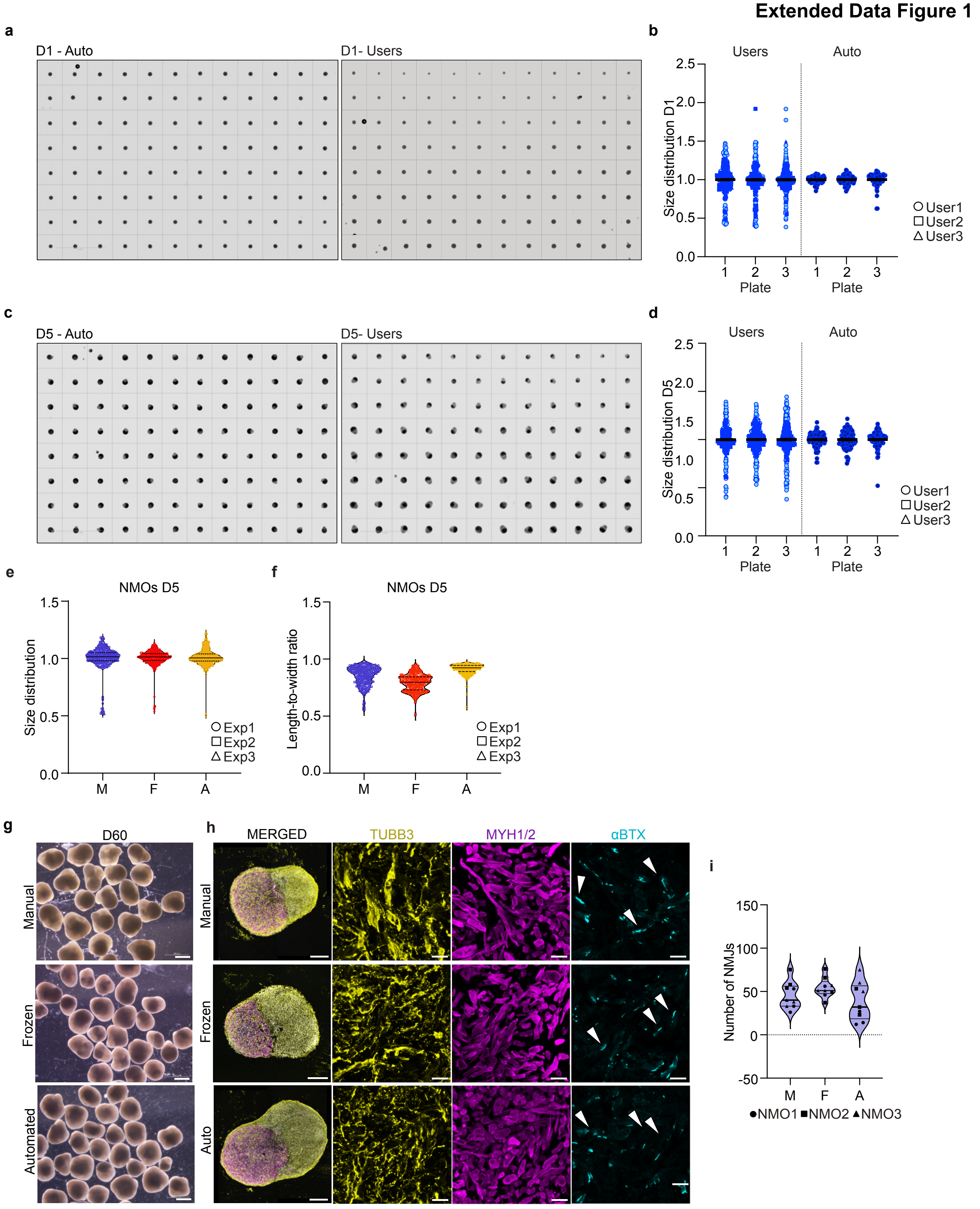

**Extended Data Figure 2**

**a**

| Cell line | Official nomenclature | Cell line ID | Source | Age at sampling | SMN1–SMN2 copy number | Sex | Mutation / Clinical diagnosis |
|---|---|---|---|---|---|---|---|
| **TTN-GFP** | WTC TTNGFP; AICS0048 | AICS-0048-039 | Allen Cell Collection | 30–34 years | 2–2 | m | Healthy control |
| **H1** | WA01 | — | WiCell | Blastocyst stage | 2–2 | m | Healthy control |
| **H9** | WA09 | — | WiCell | Blastocyst stage | 2–2 | f | Healthy control |
| **SMA Pt1** | SMAE1C4 | — | Gary J. Bassell, Wilfried Rossoll Urzi et al., *Nat Comm*, 2023. | Unknown | 0–2 | m | Early-Stage SMA Type I Pathology<br>• Presented at 8 weeks of age<br>• Acute respiratory distress and hypotonia<br>• Diagnosis consistent with Spinal Muscular Atrophy (SMA) Type I |
| **SMA Pt2** | CS84iSMA-n12 B | GM10684 Coriell | Cedars-Sinai Biomanufacturing Center | 6 months | 0–2 | f | Later-Stage SMA Type I Pathology<br>• Sampled at 6 months of age<br>• Clinical features included severe hypotonia, respiratory distress, and muscle atrophy<br>• Diagnosis confirmed by muscle biopsy<br>• Deceased at 6 months of age |
| **SMA Pt3** | CS86iSMA-n2 | GM23686 Coriell | Cedars-Sinai Biomanufacturing Center | Unknown | 0–2 | f | • Clinically affected |

**b**

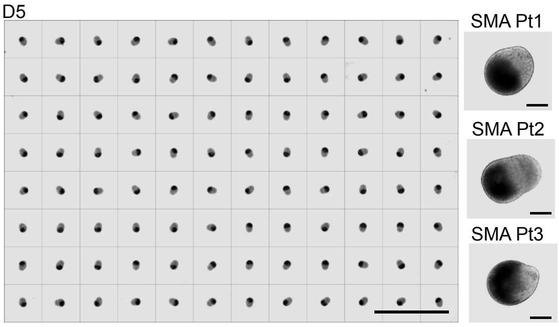


**c**

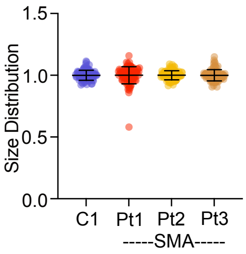


**d**

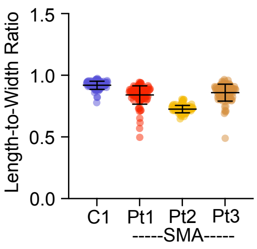

**Extended Data Figure 3**

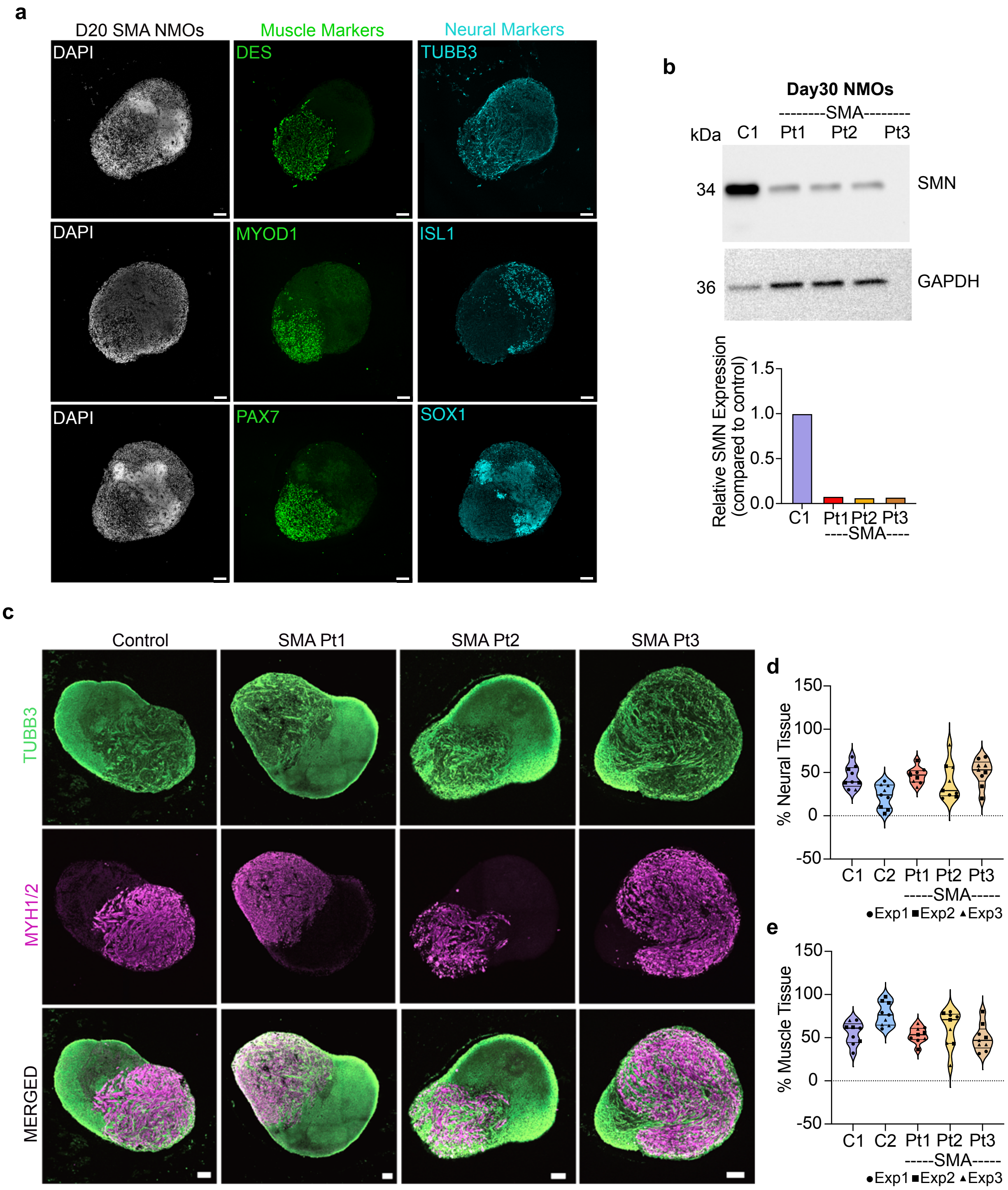

**Extended Data Figure 4**

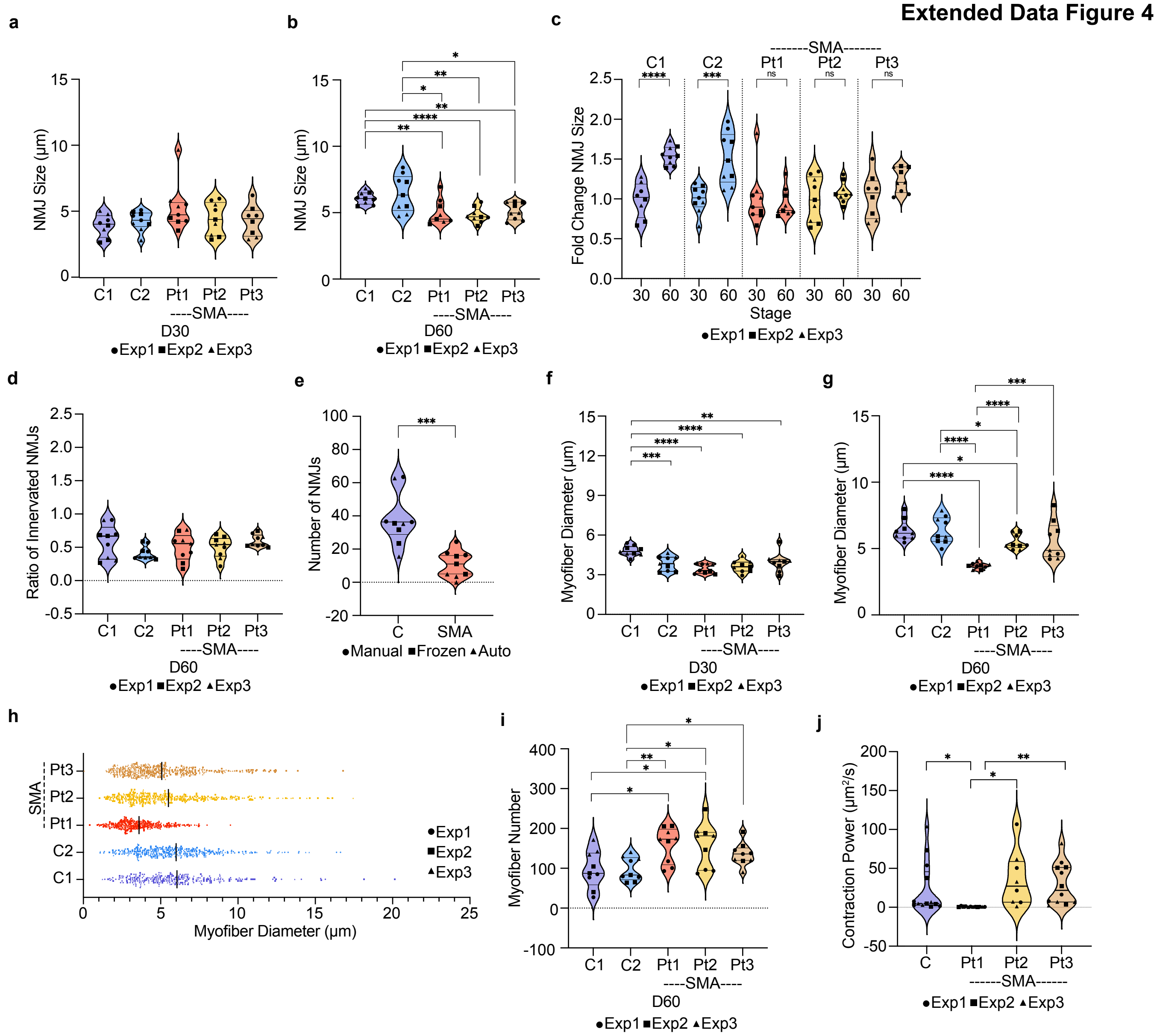

#### Extended Data Figure 5

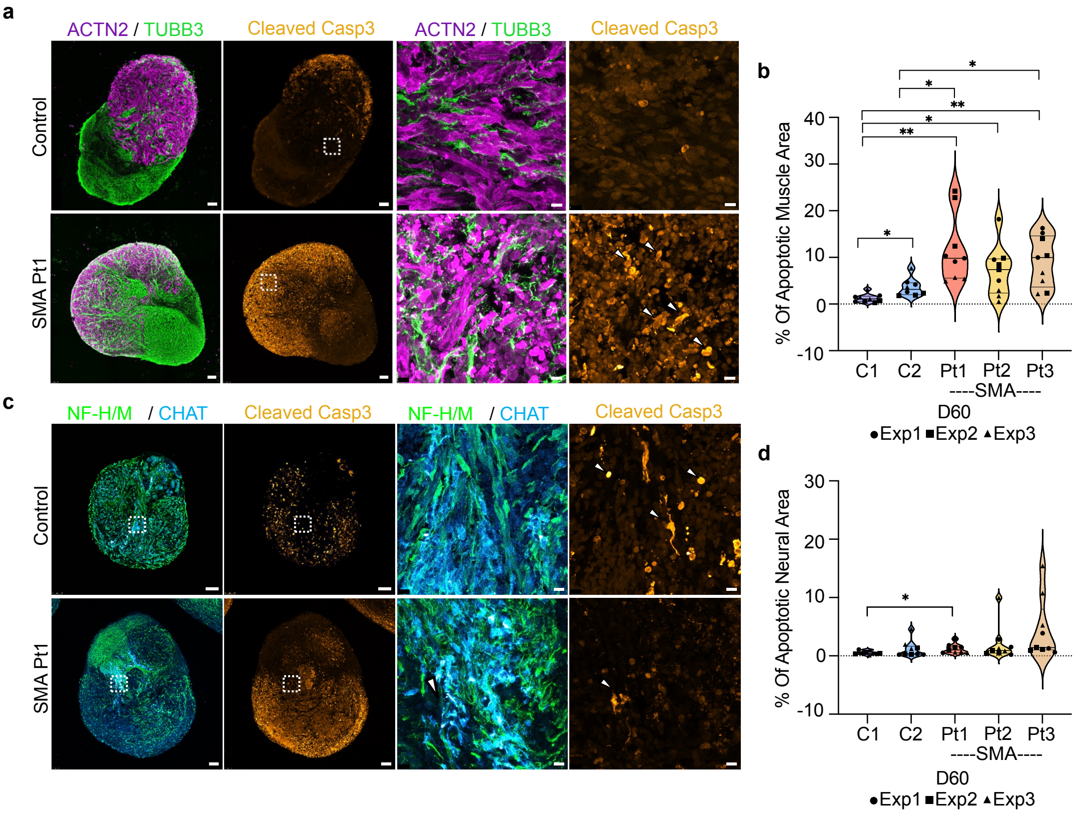

### Extended Data Figure 6

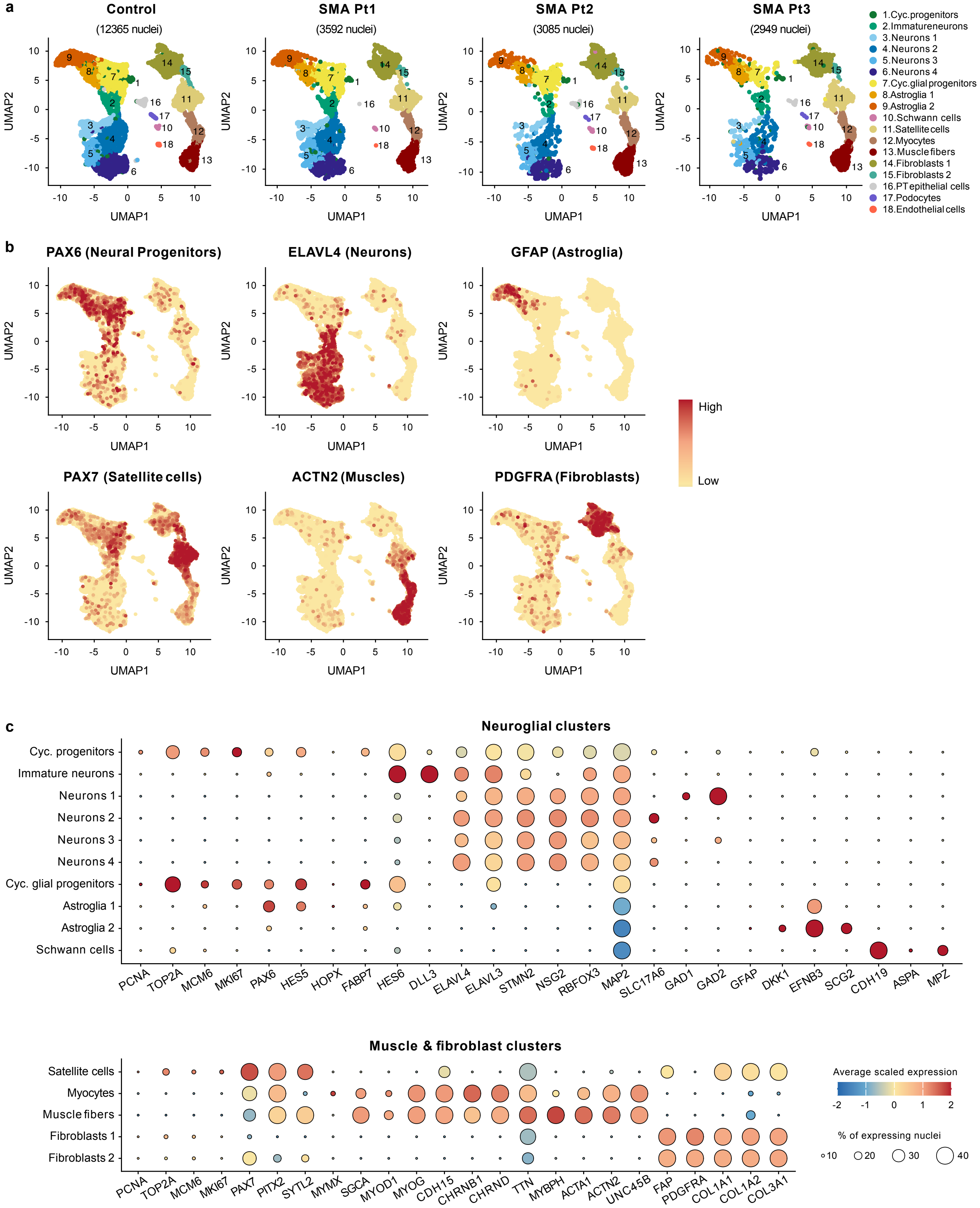

#### Extended Data Figure 7

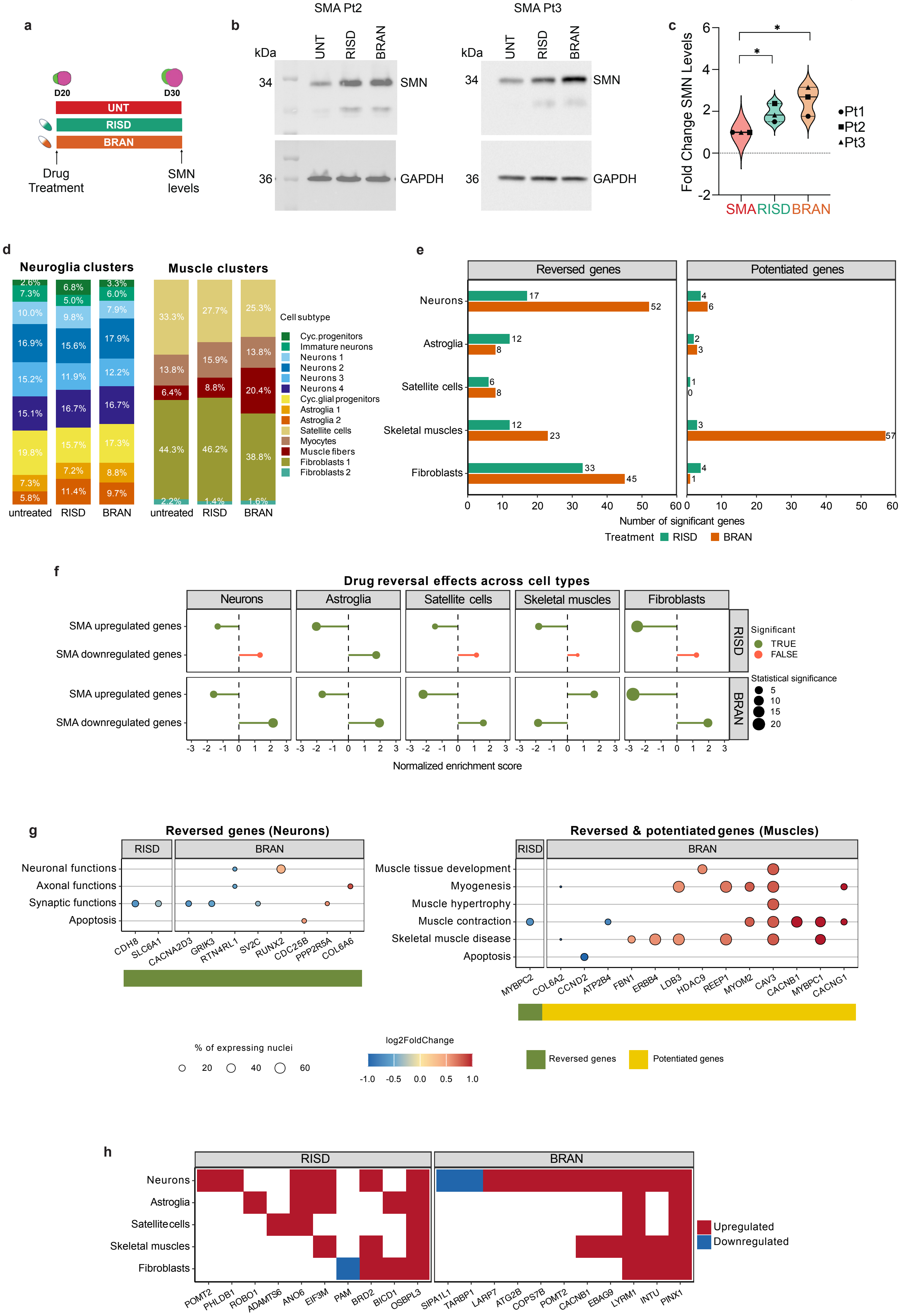

**Extended Data Figure 8**

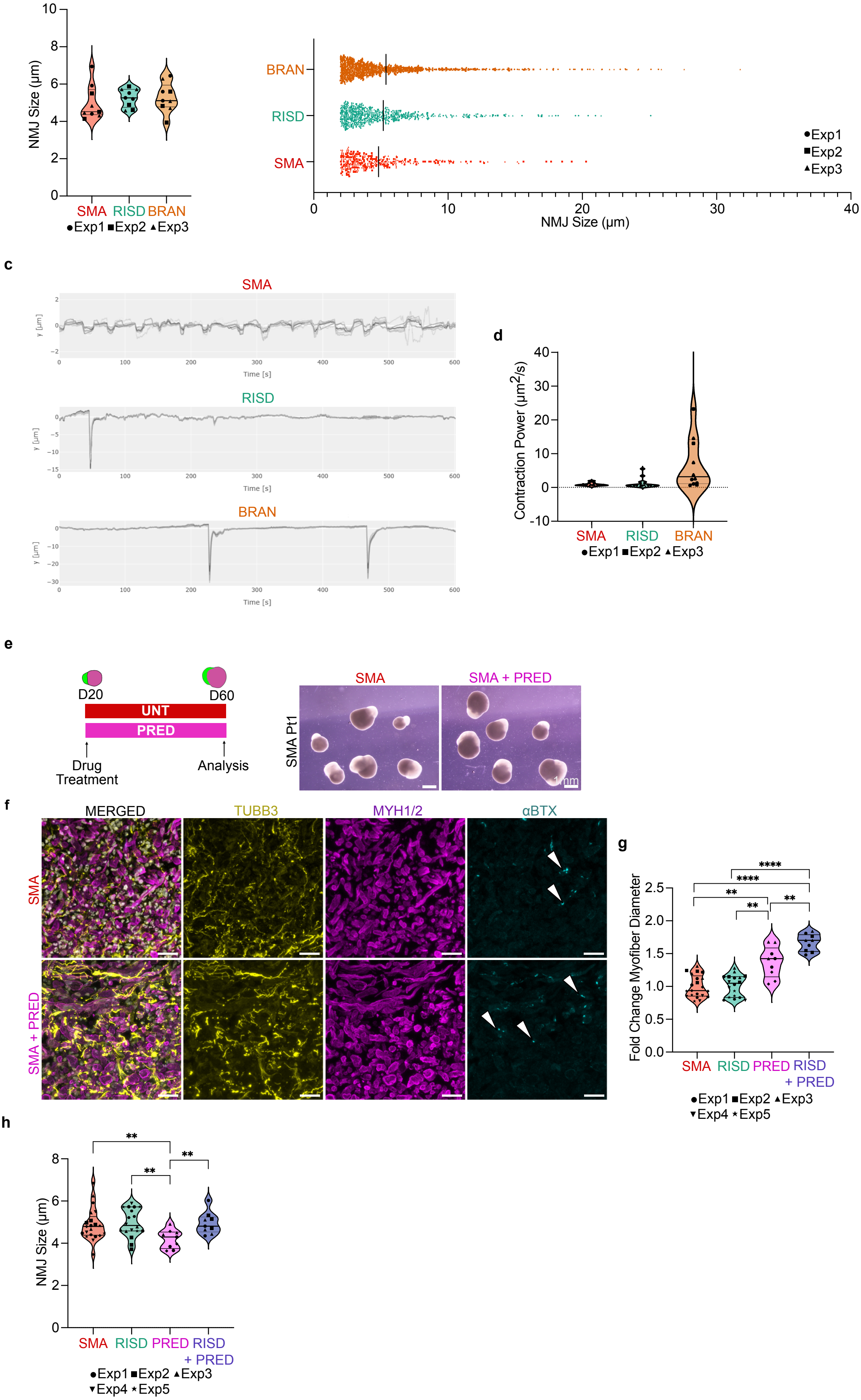
